# SpaDecon: cell-type deconvolution in spatial transcriptomics with semi-supervised learning

**DOI:** 10.1101/2023.02.12.528038

**Authors:** Kyle Coleman, Jian Hu, Amelia Schroeder, Edward B. Lee, Mingyao Li

**Affiliations:** Department of Biostatistics, Epidemiology and Informatics, Perelman School of Medicine, University of Pennsylvania, Philadelphia, PA 19104, USA; Translational Neuropathology Research Laboratory, Department of Pathology and Laboratory Medicine, Perelman School of Medicine, University of Pennsylvania, Philadelphia, PA 19104, USA

**Keywords:** cell-type deconvolution, spatial transcriptomics, histology, single-cell RNA sequencing, semisupervised learning

## Abstract

Spatially resolved transcriptomics (SRT) has advanced our understanding of the spatial patterns of gene expression, but the lack of single-cell resolution in spatial barcoding-based SRT hinders the inference of specific locations of individual cells. To determine the spatial distribution of cell types in SRT, we present SpaDecon, a semi-supervised learning approach that incorporates gene expression, spatial location, and histology information for cell-type deconvolution. SpaDecon was evaluated through analyses of four real SRT datasets using knowledge of the expected distributions of cell types. Quantitative evaluations were performed for four pseudo-SRT datasets constructed according to benchmark proportions. Using mean squared error and Jensen-Shannon divergence with the benchmark proportions as evaluation criteria, we show that SpaDecon performance surpasses that of published cell-type deconvolution methods. Given the accuracy and computational speed of SpaDecon, we anticipate it will be valuable for SRT data analysis and will facilitate the integration of genomics and digital pathology.

## Introduction

A crucial component of disease pathology is determining the relationship between cells and their relative locations. The development of spatially resolved transcriptomics (SRT) technologies has led to an increased understanding of the spatial context of gene expression in tissues [1–3]. SRT technologies identify RNA transcripts in a tissue section while specifying the approximate locations of the transcripts in the tissue. The number of genes measured and the resolution of the gene expression data depend on the type of SRT technology used to generate the data [4]. For example, fluorescence in situ hybridization (FISH)-based SRT methods, such as MERFISH [5] and seqFISH÷ [6], measure gene expression at the subcellular level, but the number of genes for which RNA transcripts can be identified is limited to at most around 10,000 [7]. On the other hand, other SRT technologies utilize spatial barcoding and next-generation sequencing techniques to measure gene expression levels in gene capture locations, referred to as spots, while retaining the relative spatial locations of spots within the tissue. Such SRT technologies include Spatial Transcriptomics (ST) [8], 10X Genomics Visium, SLIDE-seq [9], and SLIDE-seq2 [10]. While these methods provide gene expression data with lower resolution than FISH-based methods, they are generally able to measure RNA transcripts from a larger number of genes. For instance, the Visium technology measures expression levels for the entire transcriptome across spots containing between 1-10 cells on average. Also, in addition to the gene expression matrix and relative spatial locations of the spots, data from spatial barcoding-based SRT methods typically include a high-resolution hematoxylin and eosin (H&E) stained histology image of the tissue section from which the gene expression data were obtained.

When analyzing SRT data, a desirable goal is to infer the spatial locations of specific cell types. For RNA sequencing data that have single-cell resolution, a common approach is to utilize a supervised clustering algorithm that relies on cell-type specific gene expression profiles obtained from an annotated single-cell RNA sequencing (scRNA-seq) dataset. However, spatial barcoding-based SRT methods that do not have single-cell resolution can contain multiple cell types in various spots, implying that a clustering algorithm that assigns one cell type to each spot would not be adequate in localizing cell types. To understand the spatial distributions of cell types in tissues analyzed using spatial barcoding-based technologies, cell-type proportions are often estimated across the spots through a cell-type deconvolution algorithm [11].

A number of cell-type deconvolution algorithms have been developed specifically for analyzing SRT data from spatial barcoding-based technologies. RCTD [12] assumes a Poisson distribution for the count of each gene in each SRT spot and approximates cell-type proportions using maximum likelihood estimation. SPOTlight [13] is a cell-type deconvolution method for SRT data that is based on a seeded non-negative matrix factorization (NMF) regression framework. Stereoscope [14] estimates cell-type proportions using a maximum likelihood approached based on the assumption that the gene counts in the SRT and scRNA-seq data follow negative binomial distributions. Cell2location [15] utilizes a hierarchical Bayesian framework to perform cell-type deconvolution of SRT data. A major drawback of these cell-type deconvolution methods is that they do not take advantage of the high-resolution histology image and/or spatial locations of the spots for which gene expression levels are measured. This is particularly problematic since there is evidence that spots that are closer spatially and that have more similar histological features tend to have similar cell-type proportions. As a result, it can be beneficial to utilize the spatial and histological information, along with the gene expression data, to estimate cell-type proportions across spots in a tissue section analyzed using spatial barcoding-based SRT techniques.

To estimate cell-type proportions across SRT spots using gene expression, spatial location, and histology information, we developed SpaDecon, a semi-supervised learning-based method for cell-type deconvolution that can be applied to spatial barcoding-based SRT data. We analyzed eight SRT datasets, including one dataset obtained from mouse brain, three obtained from cancerous tissue sections, and four pseudo-SRT datasets constructed to evaluate the performance of SpaDecon relative to existing methods. Our results show that SpaDecon estimates cell-type proportions across spots in Visium and ST data more accurately than SPOTlight, RCTD, Stereoscope, and cell2location. Additionally, SpaDecon is computationally fast and memory efficient, making it a desirable tool for large scale SRT studies.

## Results

### Overview of SpaDecon and evaluation

An overview of the SpaDecon workflow is shown in Figure 1. Because the number of cells per spot is small (~1 to 30) [2], the observed expression at each spot is noisy. However, spots that are physically close and share similar histological features are expected to have similar gene expression and cell-type proportions, which implies that the gene expression for each spot can be denoised by aggregating information from its neighboring spots. This motivated us to use a graph-based local smoothing approach to denoise the observed gene expression.

**Fig. 1.**
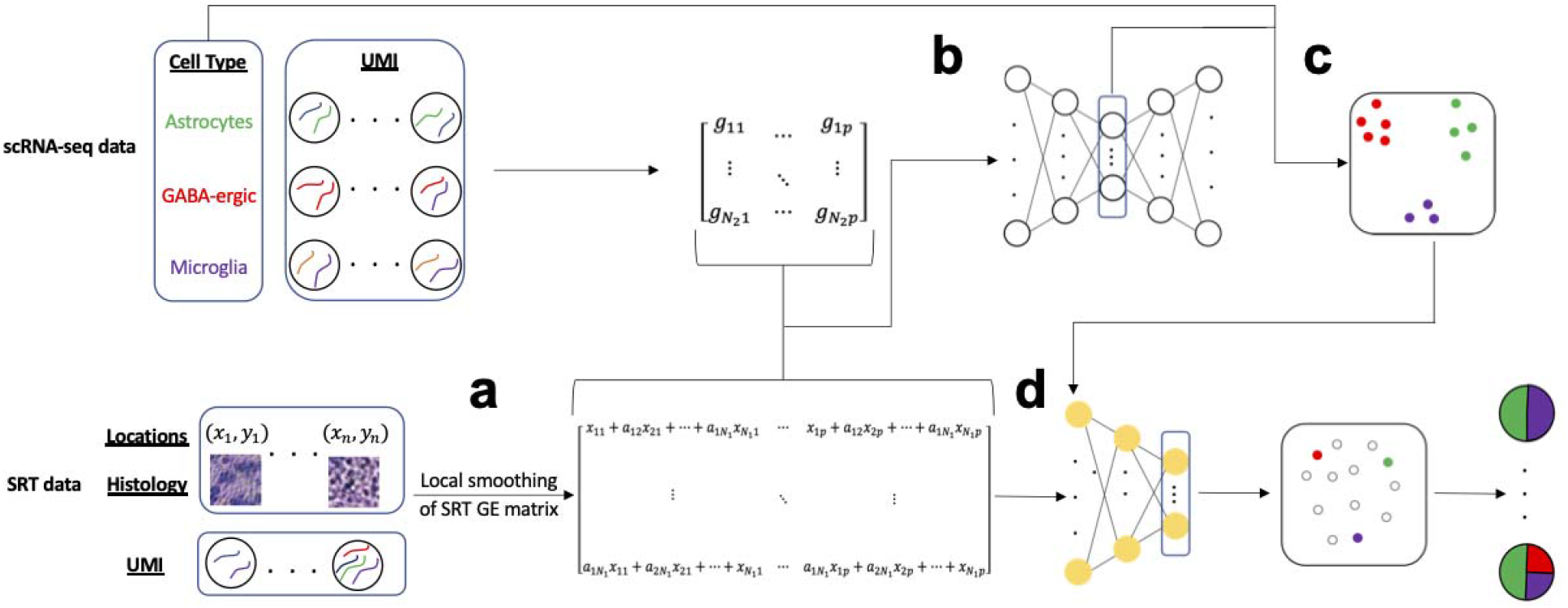
SpaDecon workflow. **a,** Using the SRT high-resolution histology image and spatial locations, SpaDecon constructs an adjacency matrix to locally smooth the SRT gene expression matrix. **b**, The smoothed SRT gene expression matrix is combined with the gene expression matrix of an annotated scRNA-seq reference dataset and used as input for a stacked autoencoder. **c,** A clustering layer is added to the final encoder layer and the annotated scRNA-seq data are used to train the network to identify features relevant for cell-type classification. **d**, The smoothed SRT gene expression matrix is used as input for the optimized network and cell-type proportions for a spot are inferred using the distances between the embedding for that spot and the centroids representing the cell types present in the scRNA-seq dataset.

SpaDecon starts by constructing an adjacency matrix that represents the similarity of spots with respect to spatial locations and histological features. Then, SpaDecon locally smooths the SRT gene expression data by performing matrix multiplication between the adjacency matrix and the SRT gene expression matrix. This local smoothing step aggregates gene expression across spots where the weight in the aggregation is determined by the physical distance and the histological feature similarity between the target spot and a neighboring spot. Using the smoothed SRT gene expression and the scRNA-seq data as input, SpaDecon then constructs a stacked autoencoder to map spots from the smoothed SRT data and cells from an annotated scRNA-seq dataset to a lower dimensional space. Once the stacked autoencoder is initialized, the decoder is removed and the remainder of the training process is driven solely by the annotated scRNA-seq data. A clustering layer is added to the final encoder layer formed by the stacked autoencoder, and an iterative cell-type clustering algorithm is used to optimize the network parameters and identify features that are relevant for cell-type classification. Each cluster centroid represents a cell type present in the scRNA-seq dataset. The smoothed SRT gene expression data are then used as input for the network and the distance between a given spot and centroid is used to estimate the proportion of cells in that spot belonging to the cell type represented by that cluster centroid.

To showcase the strength and properties of SpaDecon, we applied it to four real SRT datasets (Supplementary Table 1). We further performed benchmarking evaluations by generating SRT data with known cell-type proportions. The estimation accuracy of different cell-type deconvolution methods was evaluated by mean squared error (MSE) and Jensen-Shannon divergence (JSD) when comparing with the true cell-type proportions.

### Application to 10X Visium mouse brain serial section 1 (sagittal-anterior)

We evaluated SpaDecon’s performance on a 10X Visium dataset generated from the anterior section of a mouse brain [16]. For an annotated reference scRNA-seq dataset, we utilized the mouse whole cortex and hippocampus SMART-seq dataset from the Allen Brain Institute [17]. The original scRNA-seq dataset contained 74,973 cells classified as one of 44 cell types. To make the dataset a more suitable reference for cell-type deconvolution, we combined cells of similar types so that the scRNA-seq reference dataset contained a total of 11 cell types. Also, due to the inability of some methods, e.g., SPOTlight, to perform deconvolution with such a large reference dataset, to make a fair comparison, we sampled from this dataset to obtain a scRNA-seq reference dataset of 2,000 cells so that all methods could be run with the same single-cell reference. To evaluate SpaDecon’s performance relative to other cell-type deconvolution methods, we also ran RCTD, SPOTlight, Stereoscope, and cell2location on this dataset. Scatter pie plots and proportion heatmaps are presented in Figure 2a and Supplementary Figure 1 to show the cell-type proportions estimated by each method across all spots in this SRT dataset. In the scatter pie plot, each spot is represented as a pie chart to show the proportion of each cell type as estimated by a particular method, and the pie charts are plotted on top of the histology image.

**Fig. 2.**
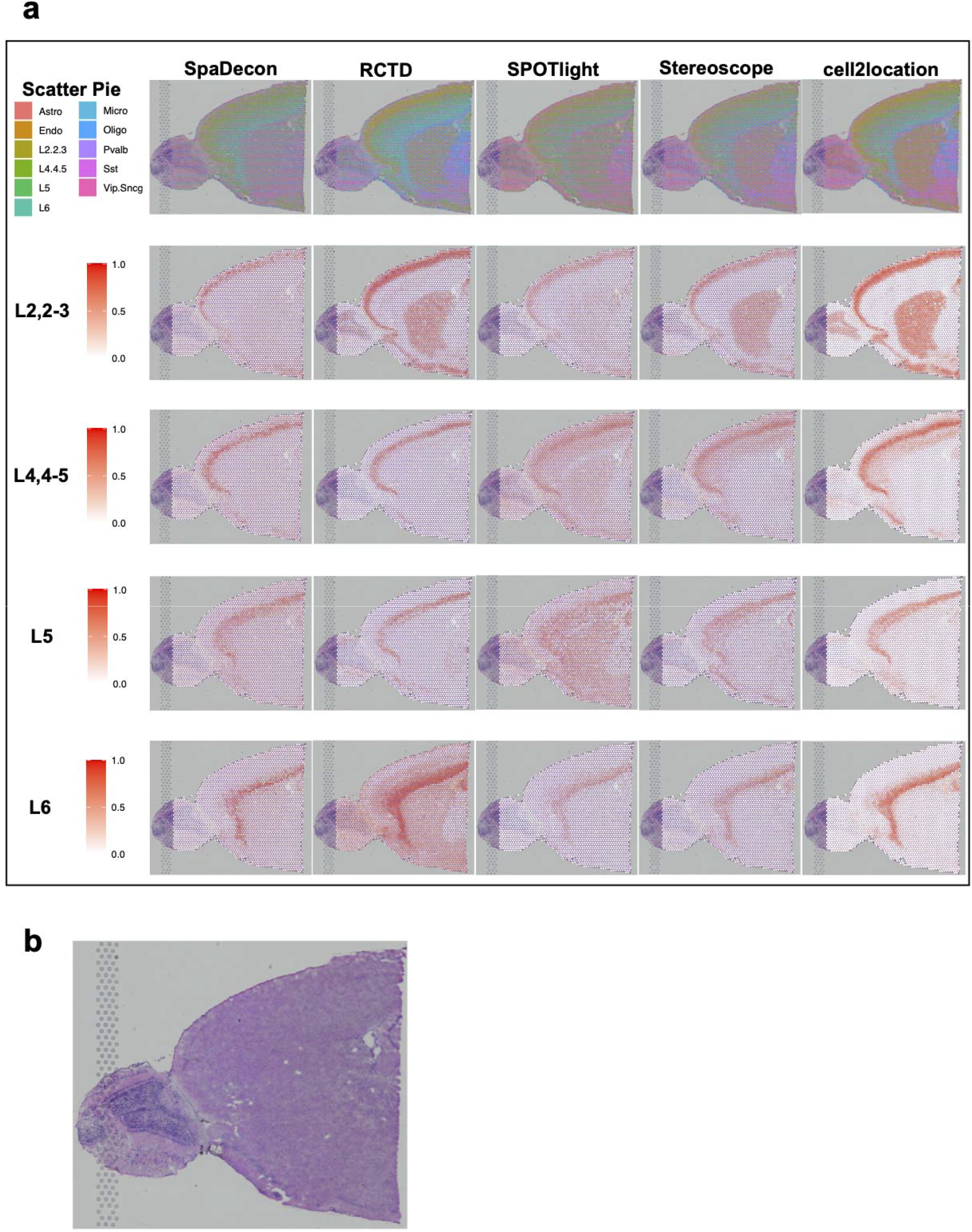
Cell-type deconvolution results for 10X Visium mouse brain serial section 1 (sagittal-anterior) data. **a,** Scatter pie plots displaying the cell-type proportion estimates of SpaDecon, RCTD, SPOTlight, Stereoscope, and cell2location at each spot, along with heatmaps showing the estimated distributions of the cortical layer-specific glutamatergic neurons. **b**, Low-resolution histology image of anterior mouse brain tissue section from which Visium data were obtained.

It is evident in the heatmaps in Figure 2a that the SpaDecon estimated proportions reveal the cortical layer structure of this anterior section of the mouse brain much more clearly than do those of the other methods. The neurons “L2,2-3”, “L4,4-5”, “L5”, and “L6” are labeled according to the cortical layers in which they are located. Hence, there should be high proportions of these cell types in their respective layers and relatively low proportions elsewhere. The heatmaps displaying the SpaDecon results agree well with this knowledge, restricting high proportions of these neurons to spots within their respective cortical layers. On the other hand, RCTD predicted high proportions of “L2,2-3” in the center of the tissue section, particularly in the caudoputamen and nucleus accumbens, and high proportions of “L6” throughout the entire tissue section. SPOTlight predicted high proportions of “L4,4-5” and “L5” throughout most of the tissue section other than the olfactory bulb. The Stereoscope and cell2location results show high proportions of “L2,2-3” in the caudoputamen and nucleus accumbens, and high proportions of “L4,4-5” throughout all cortical layers. Though we do not know the true cell-type proportions for this tissue section, the ability to localize these neurons to their proper cortical layers is evidence of SpaDecon’s superior performance compared to the other methods for this tissue section.

Since we subsampled from the single-cell reference dataset prior to running each of the methods, we constructed four additional subsamples to evaluate how this influenced the estimated proportions. For each method, we calculated the spot-level JSD between the proportions estimated using the first subsampled single-cell reference and those estimated using each of the other four subsamples. Heatmaps displaying these JSD values across all spots for each method are shown in Supplementary Figure 2. As evidenced by the heatmaps, all single-cell references resulted in similar estimated proportions for SpaDecon. The RCTD results were slightly less consistent and cell2location resulted in a large difference in proportion estimates between subsets 1 and 5. The SPOTlight and Stereoscope results were the most variable across the different single-cell datasets.

### Application to 10X Visium human breast cancer (block A section 1)

Next, we analyzed a 10X Visium dataset obtained from cancerous breast tissue [18]. We utilized an scRNA-seq reference dataset with 100,064 cells and 9 cell types [19]. We again randomly sampled 2,000 cells from this dataset so that all methods would have the ability to train on the same dataset. Heatmaps displaying the variability in the results based on the utilized single-cell subset is shown for each method in Supplementary Figure 4. Scatter pie plots and heatmaps for cancer epithelial, B, and T cells are shown in Figure 3a. Heatmaps for the 6 remaining cell types are shown in Supplementary Figure 4. The histology image is shown in Figure 3b. While all methods estimated high proportions of cancer epithelial cells throughout a majority of spots contained within the tumor regions, all methods other than SpaDecon also predicted high proportions of these cells in many non-tumor spots. This is particularly evident in the RCTD and cell2location results, where these methods estimated high proportions of cancer epithelial cells across nearly the entire tissue section. While less obvious in the SPOTlight and Stereoscope results, there are many spots in the areas adjacent to tumor regions with relatively high predicted proportions of cancer epithelial cells for these methods. SpaDecon, on the other hand, restricted high proportions of cancer epithelial cells to spots contained within the tumor regions.

**Fig. 3.**
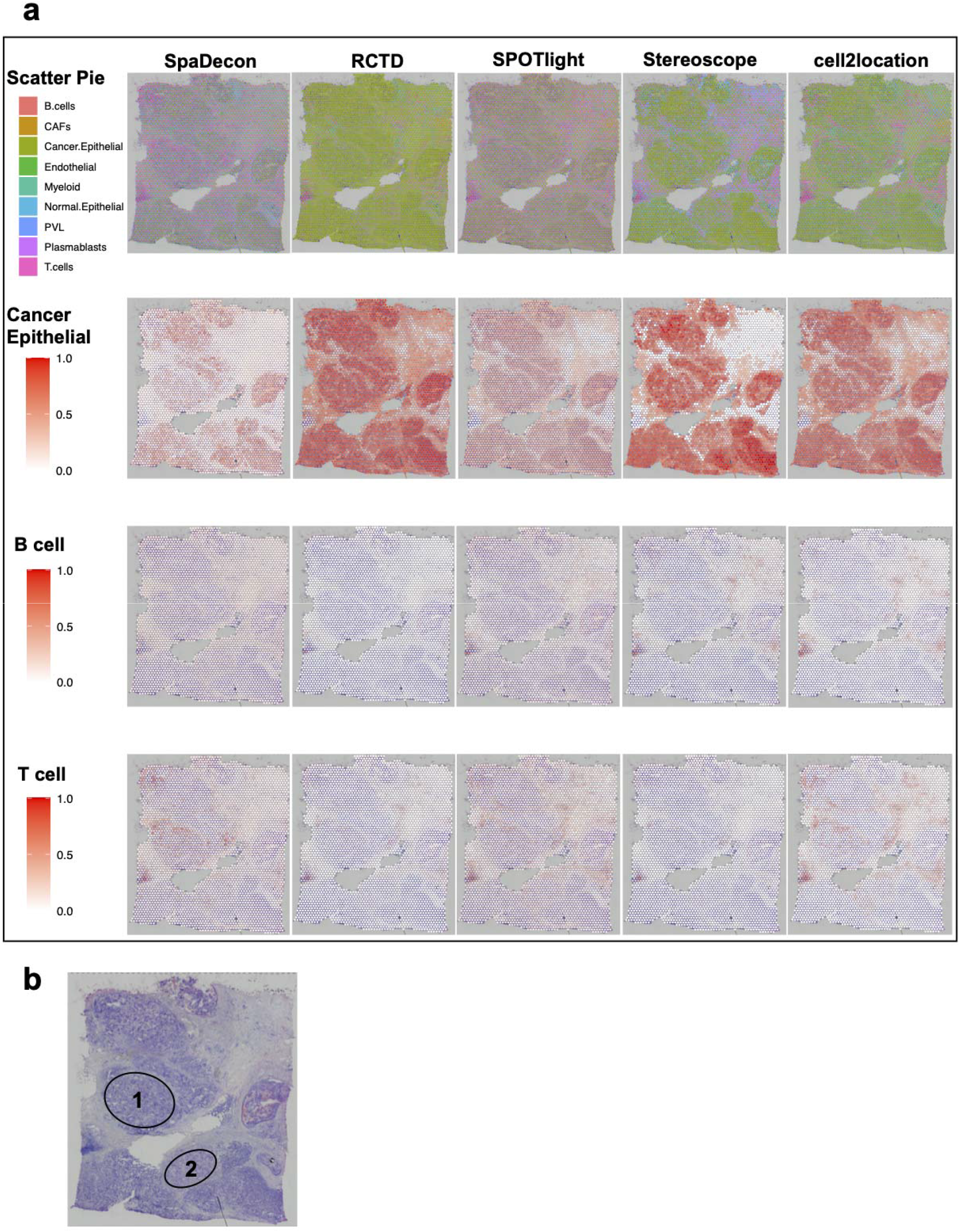
Cell-type deconvolution results for 10X Visium human breast cancer (block A section 1) data. **a,** Scatter pie plots displaying the cell-type proportion estimates of SpaDecon, RCTD, SPOTlight, Stereoscope, and cell2location at each spot, along with heatmaps showing the estimated distributions of cancer epithelial, B, and T cells. **b,** Low-resolution histology image of breast cancer tissue section from which Visium data were obtained.

There are two regions of this tissue section, annotated as “1” and “2” on the histology image, in which the SpaDecon results for cancer epithelial cells are particularly different from the other methods. For spots in these regions, RCTD, SPOTlight, Stereoscope, and cell2location predicted high proportions of cancer epithelial cells, indicating that these methods likely inferred these to be tumor regions. SpaDecon predicted high proportions of cancer epithelial cells in a few spots in these regions, but not nearly as many as the other methods. Since we are unaware of the true cell-type distributions for these data, it is not possible to definitively say which method’s results are most accurate for these regions. However, from investigation of the high-resolution histology image for these data, regions 1 and 2 have clear differences in appearance from the regions in which all methods inferred high proportions of cancer epithelial cells. This is evidence that regions 1 and 2 may not be tumor regions and contain relatively few cancer epithelial cells. If this is the case, SpaDecon likely avoided misclassifying regions 1 and 2 as tumor regions due to its ability to account for differences in histology when inferring cell-type distributions.

### Application to stage III cutaneous malignant melanoma ST data

We further utilized SpaDecon and the other cell-type deconvolution methods to analyze ST data obtained from a tissue section of a patient with stage III cutaneous malignant melanoma [20]. We used an annotated scRNA-seq dataset consisting of 7 unique cell types associated with melanoma, generated using Smart-seq 2 [21]. Scatter pie plots displaying the deconvolution results are shown in Figure 4a, along with heatmaps for malignant, B, and T cells. Heatmaps for cancer-associated fibroblasts, endothelial cells, macrophages, and natural killer cells are shown in Supplementary Figure 7. The authors that provided the ST data constructed a manual annotation of the histology image of the tissue section from which the data were obtained (Figure 4c). This annotation separates the tissue section into melanoma, stromal, and lymphoid regions. Given that the melanoma region was annotated according to where malignant cells were localized, it is expected that the proportion of this cell type in the stromal and lymphoid regions is relatively low. We obtained all spots located in the stroma and investigated the proportion of malignant cells in each spot as inferred by each deconvolution method. A boxplot displaying the proportions of malignant cells in spots contained in the stroma as estimated by each method is shown in Figure 4b. The mean proportion of malignant cells across the stromal spots as predicted by SpaDecon was 0.109, significantly less than that predicted by RCTD (mean = 0.205, p-value = 0.0003), SPOTlight (mean = 0.152, p-value = 0.03), Stereoscope (mean = 0.512, p-value = 1.57e-13), and cell2location (mean = 0.485, p-value = 4.27e-13), where the p-values were obtained using one-sided paired t-tests. This is evidence that SpaDecon more accurately localized malignant cells in this tissue section compared to the other methods. To evaluate the effect of incorporating spatial location and histology information in cell-type deconvolution, we also estimated cell-type proportions for this dataset using SpaDecon without the local smoothing step. As determined by a two-sided paired t-test, the mean proportion of malignant cells in stromal spots as predicted by SpaDecon without local smoothing was 0.105, not significantly different from that predicted by SpaDecon with local smoothing (p-value = 0.86). Hence, in this case, there is no evidence of a significant effect of incorporating histology and spatial locations through local smoothing of the gene expression matrix.

**Fig. 4.**
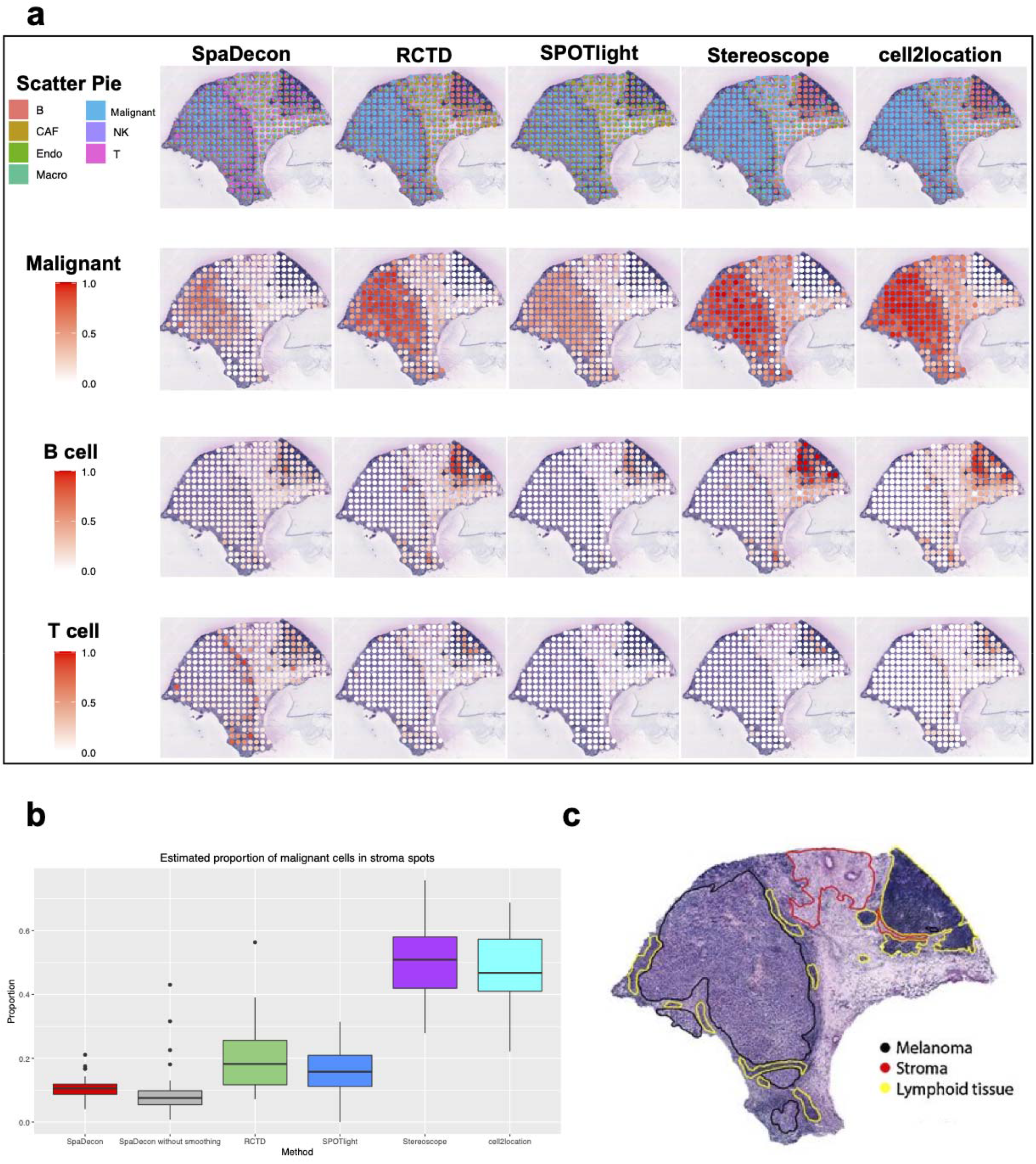
Cell-type deconvolution results for stage III cutaneous malignant melanoma ST data. **a,** Scatter pie plots displaying the cell-type proportion estimates of SpaDecon, RCTD, SPOTlight, Stereoscope, and cell2location at each spot, along with heatmaps showing the estimated distributions of malignant, B, and T cells. **b,** Boxplot showing the proportions of malignant cells across stromal spots estimated by each method. **c,** Manual annotation of histology image of melanoma tissue section from which ST data were obtained.

### Application to pancreatic ductal adenocarcinoma ST data

We also evaluated the performance of SpaDecon and the other deconvolution methods on a tumorous tissue section from a patient with untreated pancreatic ductal adenocarcinoma (PDAC), referred to as PDAC-B, analyzed using the ST technology [22]. Along with the ST data, paired scRNA-seq data were also obtained from the tumor of this patient. The scRNA-seq data consist of 13 cell types, including three unique ductal subtypes. We utilized this scRNA-seq dataset as the reference for cell-type deconvolution of the PDAC-B ST data. Annotation of the histology image, shown in Figure 5c, was provided by the authors of the paper that presented the data and reveals three distinct categories of tissue, including cancer cell-rich regions, duct epithelium, and interstitium. The cell-type deconvolution results of each method are shown in Figure 5a in the form of scatter pie plots. Heatmaps displaying the estimated distributions of cancer clone A, ductal-MHC Class II, and ductal-terminal ductal like cells are shown in Figure 5a, and heatmaps for the remaining cell types are shown in Supplementary Figure 9. The SpaDecon estimated proportions align well with the manual annotation given the cell types known to exist in each region. Specifically, SpaDecon estimated high proportions of cancer cells in the cancer cell-rich region. To quantitatively compare with the other methods, we isolated all spots contained within the annotated cancer cell-rich region. The mean SpaDecon estimated proportion of cancer cells in these spots was 0.324, higher than that estimated by RCTD (mean = 0.293, p-value = 0.20), SPOTlight (mean = 0.158, p-value = 1.40e-05), Stereoscope (mean = 0.228, p-value = 0.005), and cell2location (mean = 0.314, p-value = 0.40). We also investigated the effect of the local smoothing step of SpaDecon on the PDAC-B results. The mean proportion of cancer cells in these spots as predicted by SpaDecon without local smoothing was 0.187, significantly less than that predicted by SpaDecon with local smoothing (p-value = 0.0002). This is evidence that incorporating spatial location and histology information aids in the detection of cancer cells for this tissue section. All p-values discussed in this section were obtained using one-sided paired t-tests. A boxplot showing the proportions of cancer cells estimated by each method in cancer cell-rich region spots is shown in Figure 5b.

**Fig. 5.**
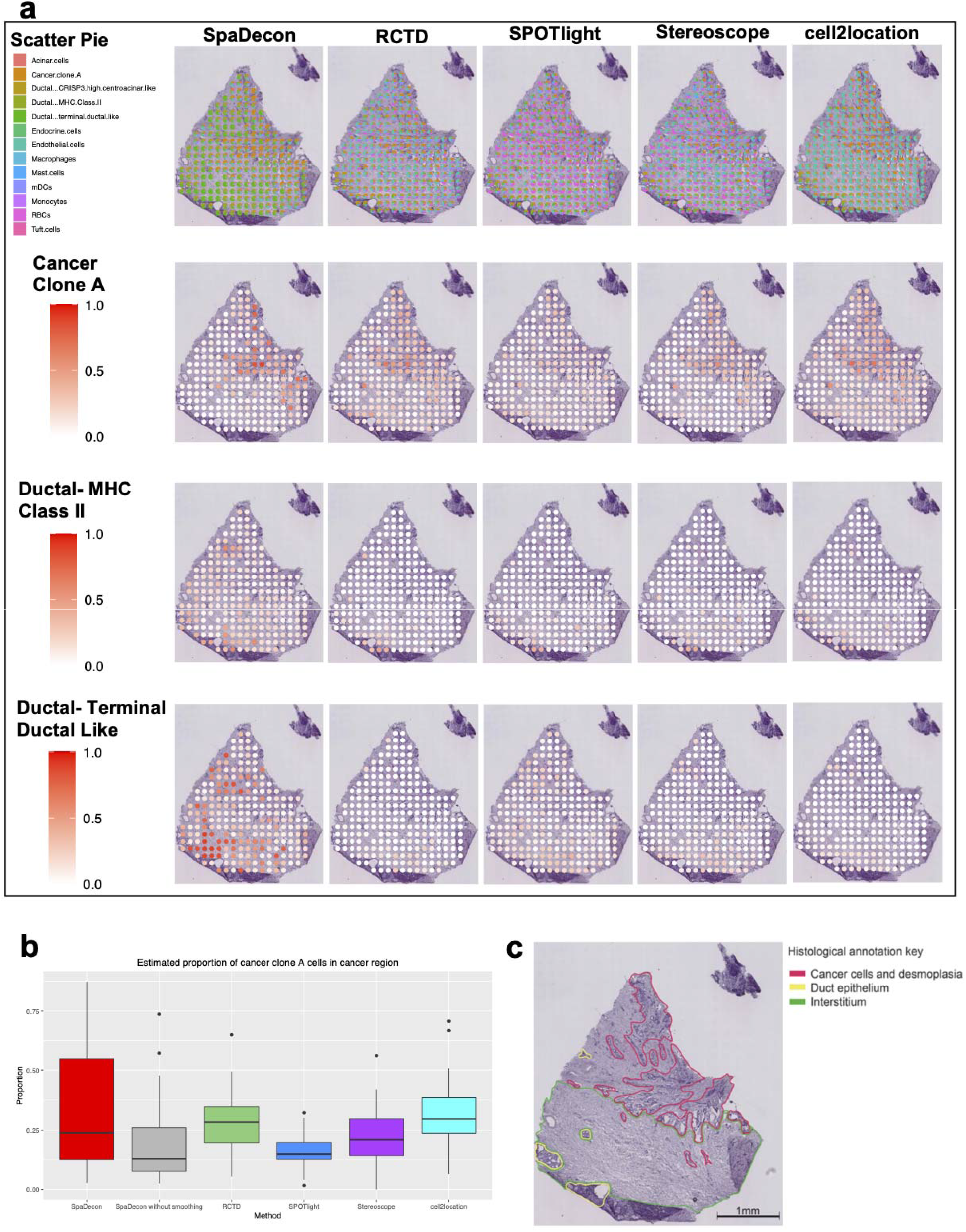
Cell-type deconvolution results for pancreatic ductal adenocarcinoma ST data. **a,** Scatter pie plots displaying the cell-type proportion estimates of SpaDecon, RCTD, SPOTlight, Stereoscope, and cell2location at each spot, along with heatmaps showing the estimated distributions of cancer clone A, ductal-MHC Class II, and ductal-terminal ductal like cells. **b**, Boxplot showing the proportions of cancer cells estimated by each method across spots in the cancer cell-rich regions. **c**, Manual annotation of cancerous pancreas tissue section from which ST data were obtained.

RCTD, SPOTlight, and Stereoscope estimated high proportions of RBCs in many of the spots throughout the tissue section. Specifically, of the 224 spots in this ST dataset, the RCTD, SPOTlight, and Stereoscope results had proportions greater than 0.1 of RBCs in 110 (49.1%), 152 (67.9%), and 165 (73.7%) spots, respectively. For 28 (12.5%) spots, SPOTlight estimated the proportion of RBCs to be greater than 0.5. Given that this is pancreatic tissue, however, the number of RBCs in the tissue section is expected to be relatively low compared to other epithelial and immune cell types. In support of this knowledge, the SpaDecon results did not report a high proportion of RBCs in any spot of the tissue section. Cell2location predicted higher proportions of RBCs compared to SpaDecon, but the maximum proportion was less than 0.05.

### Benchmark evaluations

When evaluating each of the cell-type deconvolution methods on the SRT datasets described above, since the true cell-type proportions are unknown, the relative performance of each method was assessed through visual examination of the deconvolution results and general knowledge of the approximate locations of cell types. To quantitatively compare the deconvolution results across different methods, we generated a pseudo-SRT dataset according to benchmark cell-type proportions. This pseudo-SRT dataset was constructed to resemble the PDAC-B ST dataset described earlier, and details regarding the generation of this dataset can be found in Methods. In addition to the SRT cell-type deconvolution methods evaluated on the real SRT datasets, we also evaluated MuSiC [23] on this pseudo-SRT dataset. MuSiC utilizes non-negative least squares regression to estimate cell-type proportions in bulk RNA-seq data and was not developed specifically for applications to SRT data. The performance of each deconvolution method was evaluated using the mean squared error (MSE) and Jensen-Shannon divergence (JSD) between the benchmark proportions and the estimated proportions, where the JSD between distributions *X* and *Y* is defined as

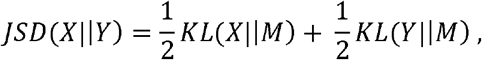

where 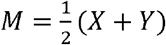 and *KL*(·) represents Kullback-Leibler (KL) divergence (defined in Methods). MSE and JSD were calculated at the cell-type level and spot level, respectively. A boxplot showing the MSE between the benchmark and estimated proportions across all cell types is shown in Figure 6a. When comparing with RCTD, SPOTlight, Stereoscope, cell2location, and MuSiC, the SpaDecon with local smoothing estimated proportions had the lowest MSE with the benchmark proportions for 6 out of 13 cell types. RCTD, SPOTlight, Stereoscope, cell2location, and MuSiC had the lowest MSE for 1, 0, 0, 2, and 4 cell types, respectively. The mean and median MSE between the SpaDecon and benchmark proportions across all cell types were 0.014 and 0.004, respectively, lower than those for RCTD (mean=0.021, median=0.017), SPOTlight (mean=0.017, median=0.008), Stereoscope (mean=0.017, median=0.014), cell2location (mean=0.020, median=0.010), and MuSiC (mean=0.016, median=0.010).

**Fig. 6.**
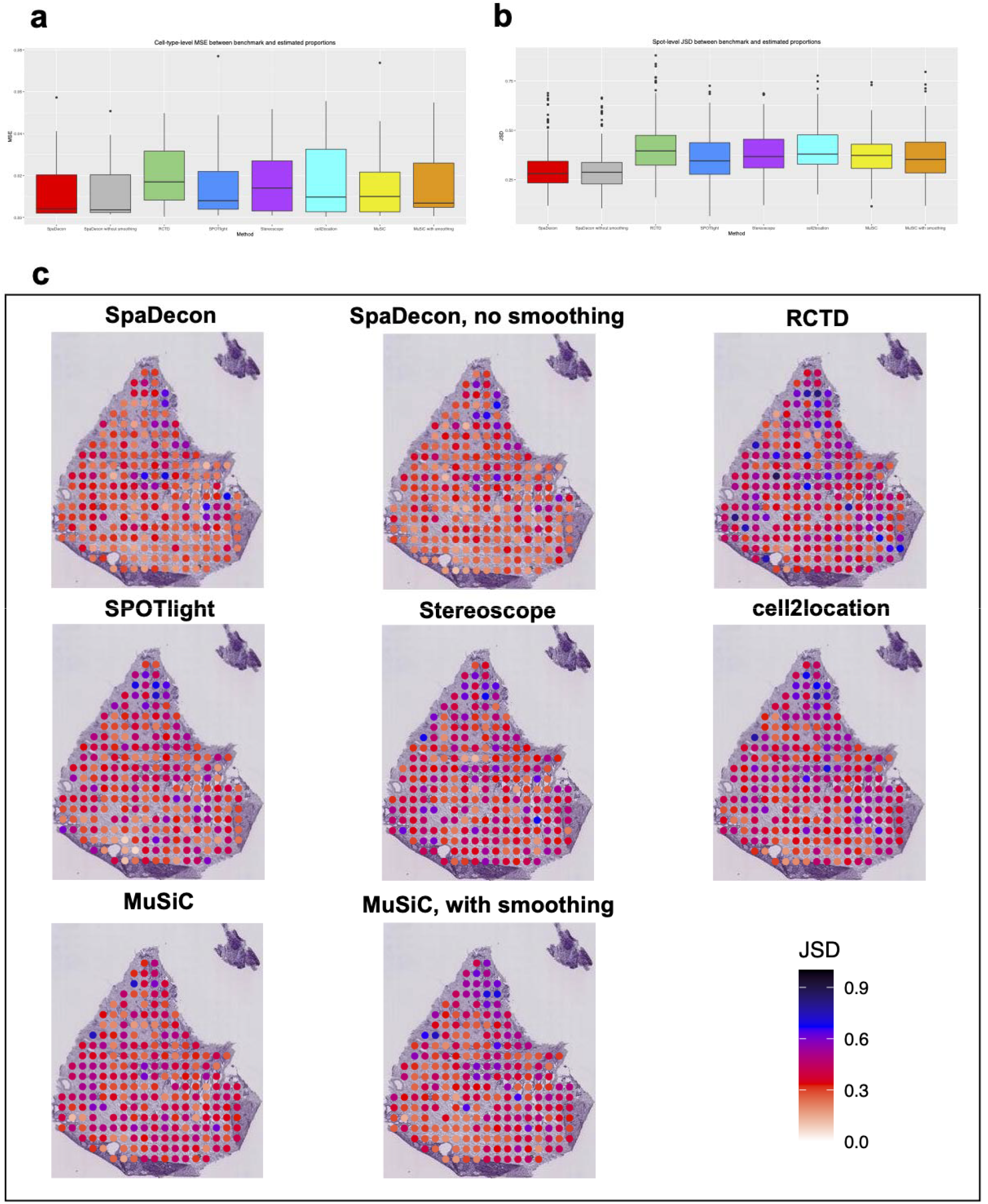
Pancreatic ductal adenocarcinoma ST benchmark evaluations. **a,** Boxplot showing the mean squared error between the benchmark and estimated proportions across all cell types for each method. **b**, Boxplot showing the Jensen-Shannon divergence between the benchmark and estimated proportions across all spots for each method. **c**, Heatmaps showing the Jensen-Shannon divergence between the benchmark and estimated proportions at each spot in the pseudo-SRT dataset for each method.

Figure 6b shows the JSD between the benchmark and estimated proportions for each method across all spots in the form of a boxplot. The mean and median JSD between the SpaDecon and benchmark proportions across all spots were 0.30 and 0.28, compared to 0.42 and 0.40 for RCTD, 0.36 and 0.35 for SPOTlight, 0.38 and 0.37 for Stereoscope, 0.40 and 0.38 for cell2location, and 0.37 and 0.37 for MuSiC. Displayed in Figure 6c are heatmaps illustrating the JSD at each spot for each method to visualize deconvolution performance in specific regions. The JSD between the SpaDecon estimated proportions and the benchmark proportions was greater than 0.5 for 15 spots, fewer than those for RCTD (44), SPOTlight (23), Stereoscope (35), cell2location (47), and MuSiC (18). Also, when comparing with RCTD, SPOTlight, Stereoscope, cell2location, and MuSiC, SpaDecon had the lowest JSD with the benchmark proportions for 104 out of 224 spots. RCTD, SPOTlight, Stereoscope, cell2location, and MuSiC had the lowest JSD for 6, 50, 30, 8, and 26 spots, respectively.

We also constructed pseudo-SRT datasets representing each of the other real SRT datasets we analyzed in this paper. The benchmark evaluation results for the 10X Visium mouse anterior brain, 10X Visium breast cancer, and stage III cutaneous malignant melanoma ST datasets are shown in Supplementary Figures 3, 6, and 8, respectively.

## Discussion

Knowledge of the spatial location information of cells within a tissue section creates many opportunities for investigating the relationships of different cells within their morphological context. While the introduction of SRT technologies has provided spatial context to gene expression data, spatial barcoding-based technologies, specifically the 10X Visium and ST, suffer from a lack of single-cell resolution. As a result, estimation of cell-type proportions enhances understanding of the distribution of cell types throughout a tissue section. In this paper we presented SpaDecon, a semi-supervised learning method for cell-type deconvolution of spatial barcoding-based SRT data that utilizes gene expression, spatial location, and histology information. Unlike many other methods developed for cell-type deconvolution in SRT, SpaDecon takes advantage of the SRT spatial locations and histology image to account for the similarity of cell-type proportions among spots that are spatially close and that have similar histological features.

Through comprehensive evaluations, we have shown that SpaDecon is effective in estimating cell-type proportions across spots in diverse Visium and ST datasets, including an anterior tissue section of the mouse brain, as well as various types of cancerous tissue. Based on visual examination and prior knowledge of the general locations of cell types, the SpaDecon estimated proportions were found to be more accurate than those of SPOTlight, RCTD, Stereoscope, and cell2location. We also quantitatively demonstrated the superior performance of SpaDecon through evaluation metrics calculated using proportions estimated for pseudo-SRT datasets. For a vast majority of cell types and spots, the SpaDecon estimated proportions were shown to have lower MSE and JSD with the benchmark proportions when compared with the other methods. Further, we provided evidence that incorporating spatial location and histology information through local smoothing in some situations lead to more accurate cell-type proportion estimates compared to using only gene expression.

In addition to utilizing informative spatial location and histology imaging data and providing accurate cell-type proportion estimates, SpaDecon is computationally fast, having analyzed 3 of the 4 real SRT datasets faster than the other four methods (Figure 7). Notably, SpaDecon has a short run time regardless of the size of the SRT dataset and the number of cell types present in the scRNA-seq reference. RCTD was fast when evaluated on ST datasets with a few hundred spots (melanoma and PDAC-B), but the run time was much longer for Visium datasets with thousands of spots (mouse anterior brain and breast cancer). Stereoscope took much longer than SpaDecon, RCTD, and SPOTlight to perform cell-type deconvolution for all SRT datasets, with a mean run time of 30 hours on a CPU. Cell2location requires a GPU to run in a reasonable amount of time and, even with a GPU, had a mean run time of over an hour. The time to train the cell2location model on a CPU can be more than 50 hours. SpaDecon’s impressive runtime can be credited, in part, to the method’s approach for dimensionality reduction of the SRT gene expression data. As detailed in Methods, SpaDecon performs highly variable gene selection and, importantly, implements a stacked autoencoder to greatly reduce the dimensionality of the data to 32-128 features per spot while maintaining the between-spot heterogeneity. The stacked autoencoder is pre-trained layer by layer, which enables us to fine-tune the entire network for a fewer number of epochs, as opposed to fully training the network with random parameter initializations. This dimensionality reduction leads to a faster runtime for the supervised version of the algorithm, in which cluster centroids are optimized in the latent feature space using the annotated scRNA-seq data. Also, SpaDecon requires a small amount of memory. Among all datasets analyzed in this paper, the maximum memory used by SpaDecon is only approximately 5.06 gigabytes (Supplementary Figure 10).

**Fig. 7.**
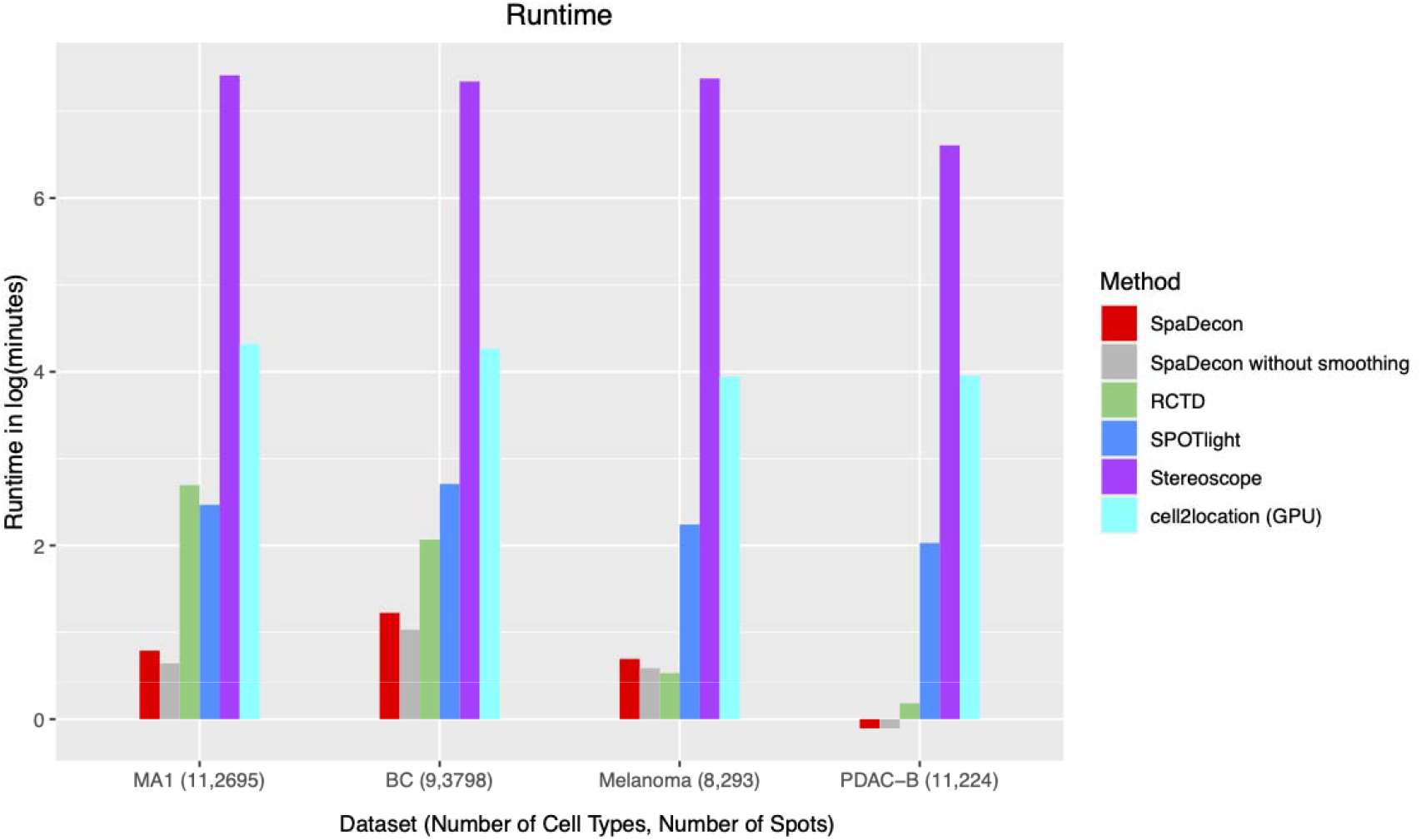
Computational run time. Bar plot displaying the run time (in log(minutes)) for each method when evaluated on each of the real SRT datasets. The number of cell types in the scRNA-seq reference datasets and the number of spots in the SRT datasets are in parentheses next to the labels of the associated SRT datasets.

In summary, through extensive evaluations both on real data and benchmark data, we have shown that SpaDecon can accurately infer the spatial distributions of cell types and is computationally fast and memory efficient. There is great potential for SRT data to provide insights into the underlying mechanisms of human disease, and we anticipate that SpaDecon will be a preferred method among biomedical researchers that seek to understand the distributions of different disease-relevant cell types throughout a tissue section.

## Methods

An overview of our cell-type deconvolution method is shown in **Fig. 7.** Because the number of cells per spot is small (~1 to 30) [2], the observed expression at each spot is noisy. However, spots that are physically close and share similar histological features are expected to have similar gene expression and cell-type proportions, which implies that the gene expression for each spot can be denoised by aggregating information from its neighboring spots.

### Input data

SpaDecon requires two datasets as input, spatial barcoding-based SRT gene expression data and annotated scRNA-seq gene expression data from the same tissue type and species. Optional input data include the 2-dimensional spatial and pixel coordinates and the histology image corresponding to the SRT gene expression data.

### Construction of adjacency matrix

When the SRT spatial locations and histology image are provided, SpaDecon utilizes these data to perform a local smoothing on the SRT gene expression matrix. To locally smooth the data, the distance between spots *u* and *v* is calculated and used to construct an adjacency matrix. The distance between *u* and *v, d*(*u,v*), accounts for the spatial location and histological features of the two spots. The spatial location for spot *v* is determined by its 2-dimensional *x* and *y* coordinates, (*x_v_,y_v_*). To calculate the difference in histology between two spots, a coordinate representing histology is constructed as in SpaGCN [24]. Specifically, the histology coordinate for spot *v, z_v_*, is defined as

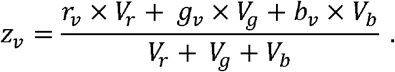

Here, *r_v_, g_v_*, and *b_v_* are the means of the RGB values across all pixels in the 50 x 50 pixel square centered on (*x_pv_,y_pv_*), where *x_pv_* and *y_pv_* are the *x* and *y* pixel coordinates for the center of spot *v*, and *V_r_, V_g_*, and *V_b_* are the variances of the RGB values across all spots in the SRT data. Defining *z_v_* in this way places the greatest weight on the RGB channel that has the largest variance across all spots. The scaled histology coordinate for spot *v* is then calculated by scaling *z_v_* such that

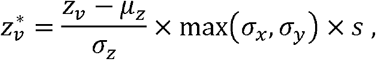

where *μ_z_* and *σ_z_* are the mean and standard deviation of *z, σ_x_* and *σ_y_* are the standard deviations of the spatial coordinates *x* and *y*, respectively, and s is the scale parameter that determines the amount of weight given to the histology data. When *s* = 1, the variance of the histology coordinates matches that of the spatial coordinate with the largest variance, giving approximately equal weight to spatial location and histology information. The coordinates of spot *v* in 3-dimensional space are then represented as 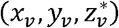, and *d*(*u, v*) is the Euclidean distance between spots *u* and *v*, i.e.,

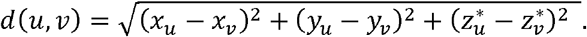

Finally, the adjacency matrix for the SRT data is constructed as ***A*** = [*a*(*u, v*)], where *a*(*u, v*), a measure of similarity between spots *u* and *v*, is defined as

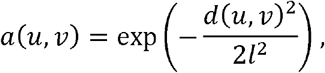

where *l* is the characteristic length scale parameter [25]. Let *X* denote the *N*_1_× *M* SRT gene expression matrix, where *N*_1_ is the number of spots and *M* is the number of genes. The smoothed gene expression matrix, *X*\*, is then constructed by performing matrix multiplication between *A* and *X*, i.e.,

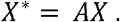

Let *A*\* = [*a*\*(*u, v*)] be defined as *A* – *I*_*N*_1__, where *I*_*N*1_ is the *N*_1_ x *N*_1_ identity matrix. So,

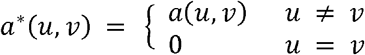

Define *a*_v_*. by:

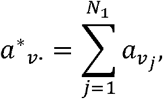

where *a*_v_j__*, the *j^th^* entry of the row of *A*\* corresponding to spot *v*, represents the weight given to the gene expression in spot *j* when calculating the smoothed expression in spot *v*. Thus, *a*_v_*. is the total weight given to the expression levels in spots *j* = 1,2, …,*v* – 1,*v* + 1, …*N*_1_ to calculate 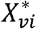 for genes *i* = 1,2, …,*M*. Let 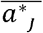. denote the mean of *a*\*_*j*_. across all spots *j* = 1, …,*N*_1_. The characteristic length scale parameter *l* is calculated automatically in SpaDecon to obtain a prespecified value of 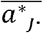. When 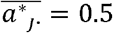, as is the default, the total weight given to the gene expression in neighboring spots is, on average, half of the weight given to the gene expression in a particular spot when calculating the smoothed expression levels for genes in that spot.

### Deep learning framework

#### Preprocessing of data

SpaDecon finds the top *h* highly variable genes (HVGs) in the smoothed SRT gene expression dataset, *X*\*, where the choice of *h* depends on the total number of genes in the dataset. Once these *h* HVGs are selected, only the *p* of these genes that are also present in the scRNA-seq gene expression dataset, *G*, are utilized. SpaDecon selects the *p* columns corresponding to these genes from *X*\* and *G*. These matrices are then logarithmized and Z-score normalized. The preprocessed SRT and scRNA-seq gene expression matrices are denoted by 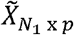 and 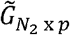, respectively.

#### Unsupervised initialization of SpaDecon model parameters

To initialize the weights of the model, SpaDecon combines the preprocessed scRNA-seq and SRT data to form the matrix *Y*_(*N*_1_+*N*_2_) *xp*_ and searches for a mapping to a lower dimensional space, 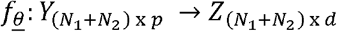 where 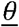 is a vector of embedding parameters and *d* « *p*. This mapping is identified using a stacked autoencoder, in which each layer is a denoising autoencoder and the number of layers depends on the number of cells and spots in the scRNA-seq and SRT datasets, respectively. The SRT data are unlabeled, and the scRNA-seq cell-type labels are not used for initialization of the stacked autoencoder, so this step is fully unsupervised. The layers of the stacked autoencoder are trained sequentially. For a given layer of the stacked autoencoder, the input is a corrupted version of the output of the previous layer, in which the values for some features are randomly set to 0. The autoencoder is then trained to reconstruct the uncorrupted features of the previous layer by minimizing the mean squared error (MSE) between the uncorrupted input and the reconstruction. For all layers in the stacked autoencoder other than the bottleneck and final decoder layers, ReLU is used as the activation function. For the bottleneck and final decoder layers, however, tanh is used as the activation since the features in these layers must have the ability to take on positive and negative values. Once each layer is trained, SpaDecon combines all encoders in layer-wise training order, followed by all decoders in reverse layer-wise training order. Combining the encoders and decoders in such a way produces a stacked autoencoder with a bottleneck layer. Once the parameters of the stacked autoencoder are fine-tuned by minimizing the MSE between the original features and the output of the final decoder layer, the decoder layers are removed. The mapping from the original features to the features in the bottleneck layer is the desired initial mapping to the latent feature space, 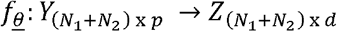.

#### Supervised optimization of SpaDecon model parameters

Once the network parameters 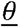 are initialized, the SRT data are removed from the training process, and the labeled scRNA-seq data are used to fine-tune the weights in a supervised fashion. We add a clustering layer that is fully connected to the final encoder layer of the stacked autoencoder described previously. Each cell type is represented by a centroid in the clustering layer, and the weights of the model are optimized by maximizing the probability of assigning the cells to their respective types in the latent feature space. We first define the cluster centroid for cell type *j* to be the mean of the latent features of all cells of type *j*. Let *q_ij_* denote the similarity between the embedded point for cell *i*, 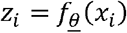, and the cluster centroid for cell type *j, μ_j_*. This similarity *q_ij_* is defined using the Student’s *t*-distribution as a kernel [26]. Specifically,

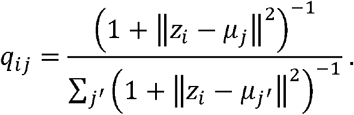

We then define a target distribution *R* as

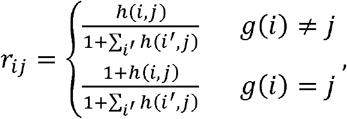

where *h*(*i,j*) is minimal random noise, *g*(*i*) is the cell type of cell *i*, and ∑_*i*_*r_ij_* = 1. The network parameters and cluster centroids are simultaneously optimized by minimizing the Kullback-Leibler (KL) divergence loss between *q_ij_* and *r_ij_* through stochastic gradient descent (SGD) with momentum, where

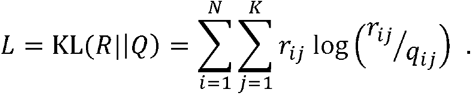

By defining the target distribution and loss function in this way, SpaDecon iteratively reduces the distance between the embedded points for the cells and the cluster centroids of their respective types. The algorithm for optimization of the network parameters and cluster centroids converges when either KL(*R*||*Q*) decreases by less than 0.01 between any two consecutive updates or the maximum number of epochs, 2000 by default, is reached.

#### Inference of cell-type proportions for SRT spots

Once the SpaDecon model has been optimized, we use the smoothed and preprocessed SRT GE matrix 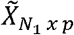 as input to the network. For spot *i* and cell type *j*, the output of the clustering layer described above, *q_ij_*, is a measure of similarity between *z_i_*, i.e., the embedding of spot *i*, and *μ_j_*, i.e., the cluster centroid for cell type *j*. Following the Deep Embedded Clustering paper [27], *q_ij_* is interpreted as the probability of assigning spot *i* to cell type *j*, which is an approximate of the proportion of cells in spot *i* belong to cell type *j*. Similar cell type proportion estimation procedure has been utilized by DSTG [28], which calculates the relative distances of spots to pseudo-spots of known composition, but in SpaDecon, we use the centroids of purely one cell type as the centroid when estimating the cell type proportions.

### Construction of pseudo-SRT data for benchmark evaluations

The pseudo-SRT dataset discussed in the results section was constructed to accurately resemble the true SRT gene expression matrix for the PDAC-B ST dataset. Specifically, the total read count of each spot in the pseudo-SRT data is equal to the total unique molecular identifier (UMI) count of the corresponding spot in the ST data. For each spot, we sampled cells from the paired scRNA-seq according to benchmark proportions. A goal for our benchmark evaluations was to quantify each method’s ability to localize rare cell types. Thus, to construct the benchmark proportions, we first classified two cell types as rare cell types. These rare cell types, determined to be macrophages and tuft cells, had the smallest mean proportions across all spots of the PDAC-B dataset when averaging across the results of SpaDecon, RCTD, SPOTlight, Stereoscope, and cell2location. For each spot, we set the benchmark proportions for these rare cell types equal to the minimum proportions estimated for that cell type across for all methods for that spot. For all other cell types, the benchmark cell-type proportions for a given spot were set equal to the mean of the cell-type proportions estimated by SpaDecon, RCTD, SPOTlight, Stereoscope, and cell2location for that spot. We then normalized the benchmark proportions so that they would sum to 1 for each spot. Generating the pseudo-SRT data in this way ensured that the evaluation metrics would reveal performance quality on realistic SRT data. Since the UMI counts differ in each spot of the SRT data, we sampled a different number of cells for each spot of the pseudo-SRT dataset. The scRNA-seq gene expression data were provided in transcripts per million (TPM) format, so the expression measures for all genes for a cell sum to approximately one million. This is far greater than the total UMI count for a given spot, as the mean total UMI count across all spots is approximately 2400. We therefore divided the expression measures for all genes in all cells in the scRNA-seq dataset by 1000, so that the sum of counts of all genes for a cell is 1000 and so that we would sample, on average, approximately 2.4 cells per spot. The number of cells of type *j* sampled for a given spot *i* is

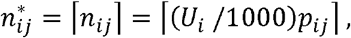

such that 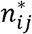 is the smallest integer greater than or equal to (*U_i_* /10000)*p_ij_*, where *U_i_* is the total UMI count for spot *i* from the SRT data and *p_ij_* is the benchmark proportion of cell type *j* in spot *i*. For the 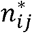 cells of type *j* that are randomly sampled, the counts for all genes in these cells are multiplied by 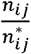 to account for sampling 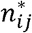 cells of type *j* instead of *n_ij_* cells. So, if *m_ijkl_* is the count for gene *k* of cell *I* sampled from cell type *j* for spot *i*, the contribution of cell *l* of type *j* to the count of gene *k* in spot *i* in the pseudo-SRT data is

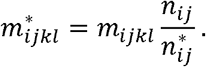

The counts of all genes are summed over all cells that are sampled for spot *i*, and so the count for gene *k* in spot *i* of the pseudo-SRT data is given by

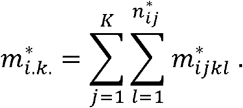

Thus, since there are 224 spots in the SRT dataset and 19738 genes in the scRNA-seq dataset, the pseudo-SRT gene expression matrix is constructed as

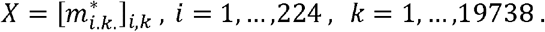

Since RCTD, SPOTlight, Stereoscope, and cell2location require a matrix of raw read counts as input for the SRT data, we rounded all values in *X* prior to evaluation of all methods. We used PDAC-B paired scRNA-seq data as the reference dataset to obtain each method’s cell-type deconvolution results for the pseudo-SRT dataset.

All other pseudo-SRT datasets were constructed analogously to the pseudo-SRT dataset described above. MuSiC was unable to analyze the rounded pseudo-SRT dataset representing the stage III cutaneous malignant melanoma dataset, and so we evaluated MuSiC on the pseudo-SRT dataset prior to rounding.

