## Supplementary Information for "SpaDecon: cell-type deconvolution in spatial transcriptomics with semi-supervised learning"

**Supplementary Table 1.** Datasets analyzed in this paper.

| Species | Tissue | Data Source | Dataset Dimensions | Protocol |
|---|---|---|---|---|
| Mouse | Anterior brain (sagittal) | 10X Genomics https://support.10xgenomics.com/spatial-gene-expression/datasets/1.1.0/V1_Mouse_Brain_Sagittal_Anterior | 2,695 spots<br>32,285 genes | 10x Visium |
| Mouse | Whole cortex and hippocampus | Yao *et al.* [1] https://portal.brain-map.org/atlases-and-data/rnaseq/mouse-whole-cortex-and-hippocampus-smart-seq | <u>Original:</u><br>76,533 cells<br>45,768 genes<br><br><u>Reduced:</u><br>2,000 cells<br>45,768 genes | SMART-seq |
| Human | Invasive Ductal Carcinoma breast tissue | 10X Genomics https://support.10xgenomics.com/spatial-gene-expression/datasets/1.1.0/V1_Breast_Cancer_Block_A_Section_1 | 3,798 spots<br>36,601 genes | 10x Visium |
| Human | Breast cancer tissue | Wu *et al.* [2]<br>GSE176078 | <u>Original:</u><br>100,064 cells<br>29,733 genes<br><br><u>Reduced:</u><br>2,000 cells<br>29,733 genes | 10x Chromium |
| Human | Stage III cutaneous malignant melanoma tissue | Thrane *et al.* [3] https://www.spatialresearch.org/resources-published-datasets/doi-10-1158-0008-5472-can-18-0747/ | 293 spots<br>16,148 genes | Spatial Transcriptomics |
| Human | Metastatic melanoma tissue | Tirosh *et al.* [4]<br>GSE72056 | 4,139 cells<br>23,686 genes | Smart-seq2 |
| Human | Pancreatic adenocarcinoma tissue | Moncada *et al.* [5]<br>GSM3405534 | 224 spots<br>19,738 genes | Spatial Transcriptomics |
| Human | Pancreatic adenocarcinoma tissue | Moncada *et al.* [5]<br>GSE111672 | 1,733 cells<br>19,738 genes | inDrop |

**Supplementary Table 2.** Software compared with SpaDecon.

| Method | Version | URL | Reference |
|---|---|---|---|
| RCTD | 1.2.0 | https://github.com/dmcable/RCTD | [6] |
| SPOTlight | 0.1.7 | https://github.com/MarcElosua/SPOTlight | [7] |
| Stereoscope | 0.2.0 | https://github.com/almaan/stereoscope | [8] |
| cell2location | 0.1.0 | https://github.com/BayraktarLab/cell2location | [9] |
| MuSiC | 0.2.0 | https://github.com/xuranw/MuSiC | [10] |

**Supplementary Table 3.** Cell type abbreviations.

| Abbreviation | Full Name |
|---|---|
| Astro | Astrocytes |
| CAFs | Cancer-associated fibroblasts |
| Endo | Endothelial cells |
| Macro | Macrophages |
| mDCs | Myeloid dendritic cells |
| Micro | Microglia |
| NK | Natural killer cells |
| Oligo | Oligodendrocytes |
| Pvalb | Parvalbumin |
| PVL | Perivascular-like cells |
| RBCs | Red blood cells |
| Sst | Somatostatin-expressing neurons |
| Vip.Sncg | Vasoactive intestinal polypeptide/ synuclein-gamma expressing neurons |

**Supplementary Figure 1.** For each method, heatmaps displaying the estimated distributions of cell types across the 10X Visium mouse anterior brain dataset. Each spot is colored according to the proportion of a given cell type in that spot as estimated by a given method. See Supplementary Table 3 for full names of cell types.

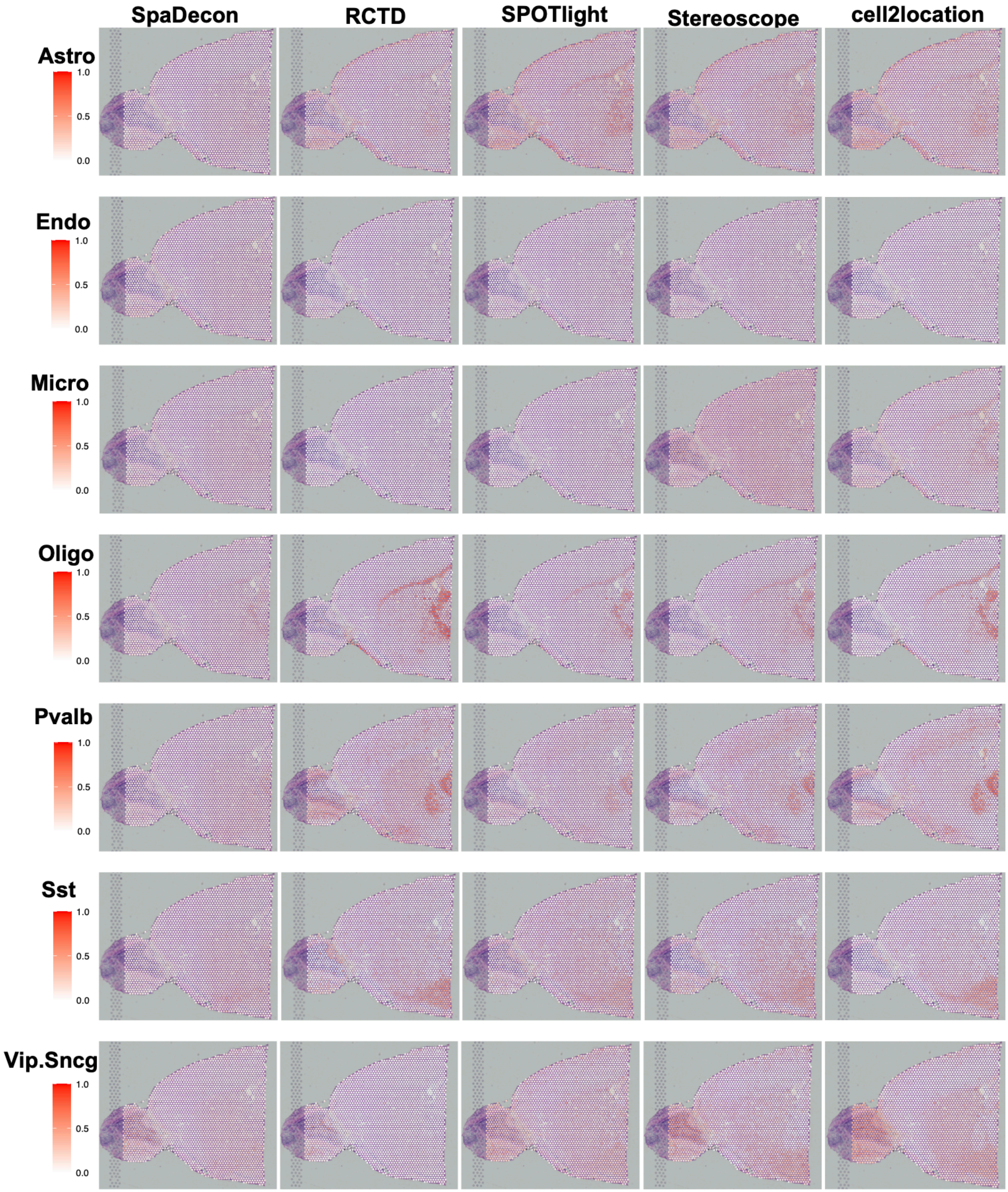

**Supplementary Figure 2.** For each method, heatmaps displaying the Jensen-Shannon Divergence (JSD) between the proportions estimated using the first scRNA-seq subset as reference and those estimated using each following scRNA-seq subset as reference for cell-type deconvolution of the 10X Visium mouse anterior brain dataset.

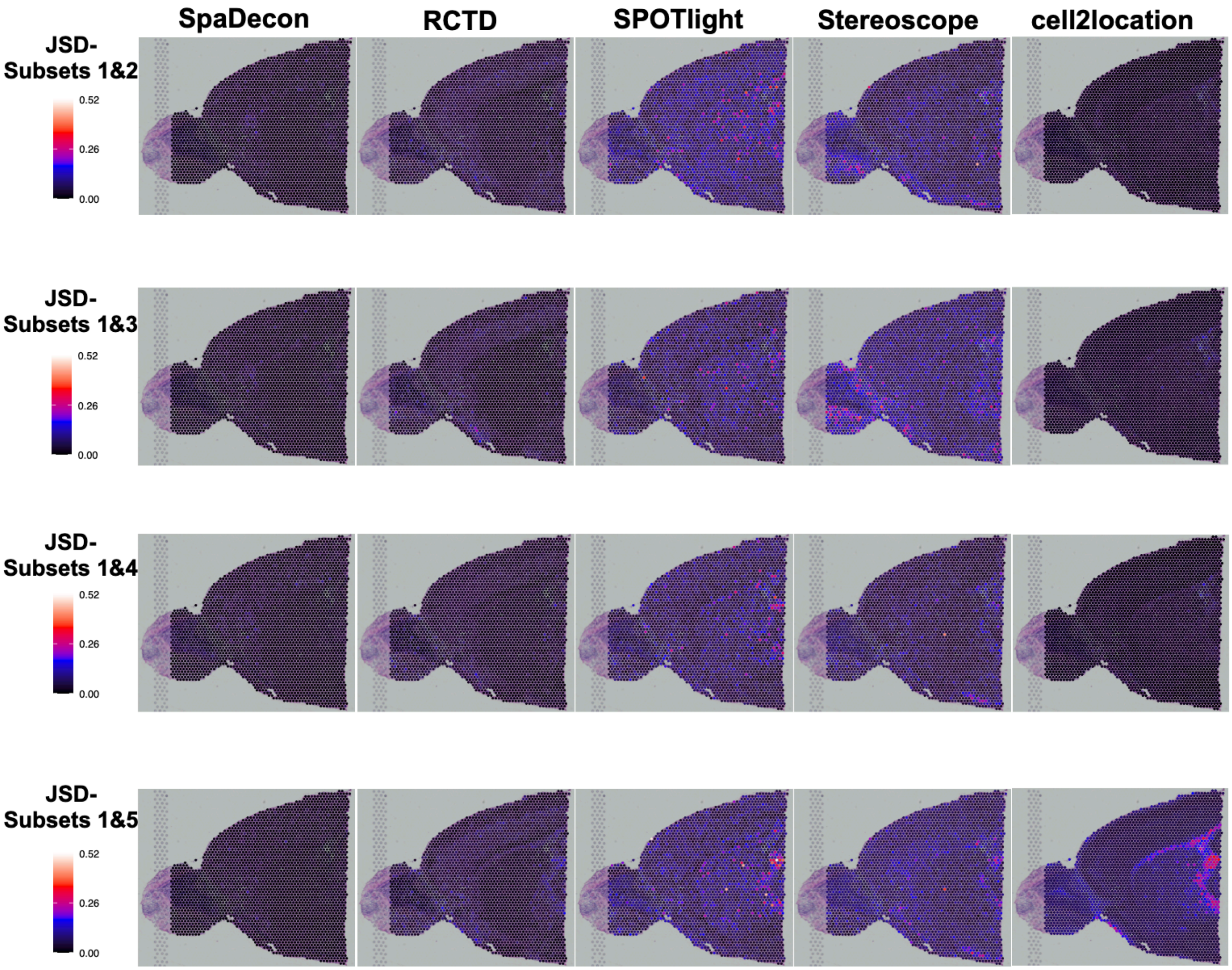

**Supplementary Figure 3. 10X Visium mouse anterior brain benchmark evaluations. a,** Boxplot showing the mean squared error between the benchmark and estimated proportions across all cell types for each method. **b,** Boxplot showing the Jensen-Shannon divergence between the benchmark and estimated proportions across all spots for each method. **c,** Heatmaps showing the Jensen-Shannon divergence between the benchmark and estimated proportions at each spot in the pseudo-SRT dataset for each method.

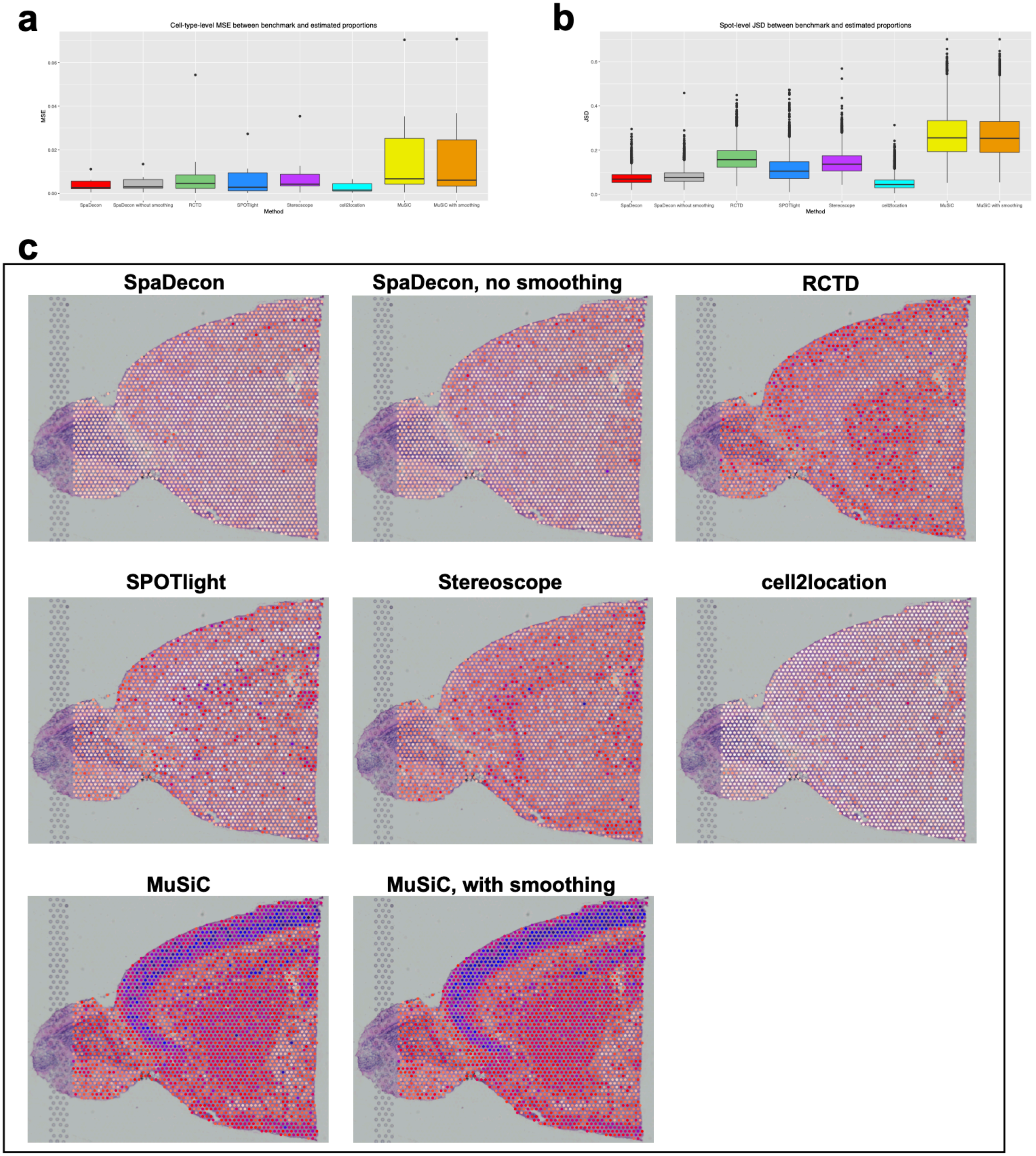

**Supplementary Figure 4.** For each method, heatmaps displaying the estimated distributions of cell types across the 10X Visium breast cancer dataset. Each spot is colored according to the proportion of a given cell type in that spot as estimated by a given method. See Supplementary Table 3 for full names of cell types.

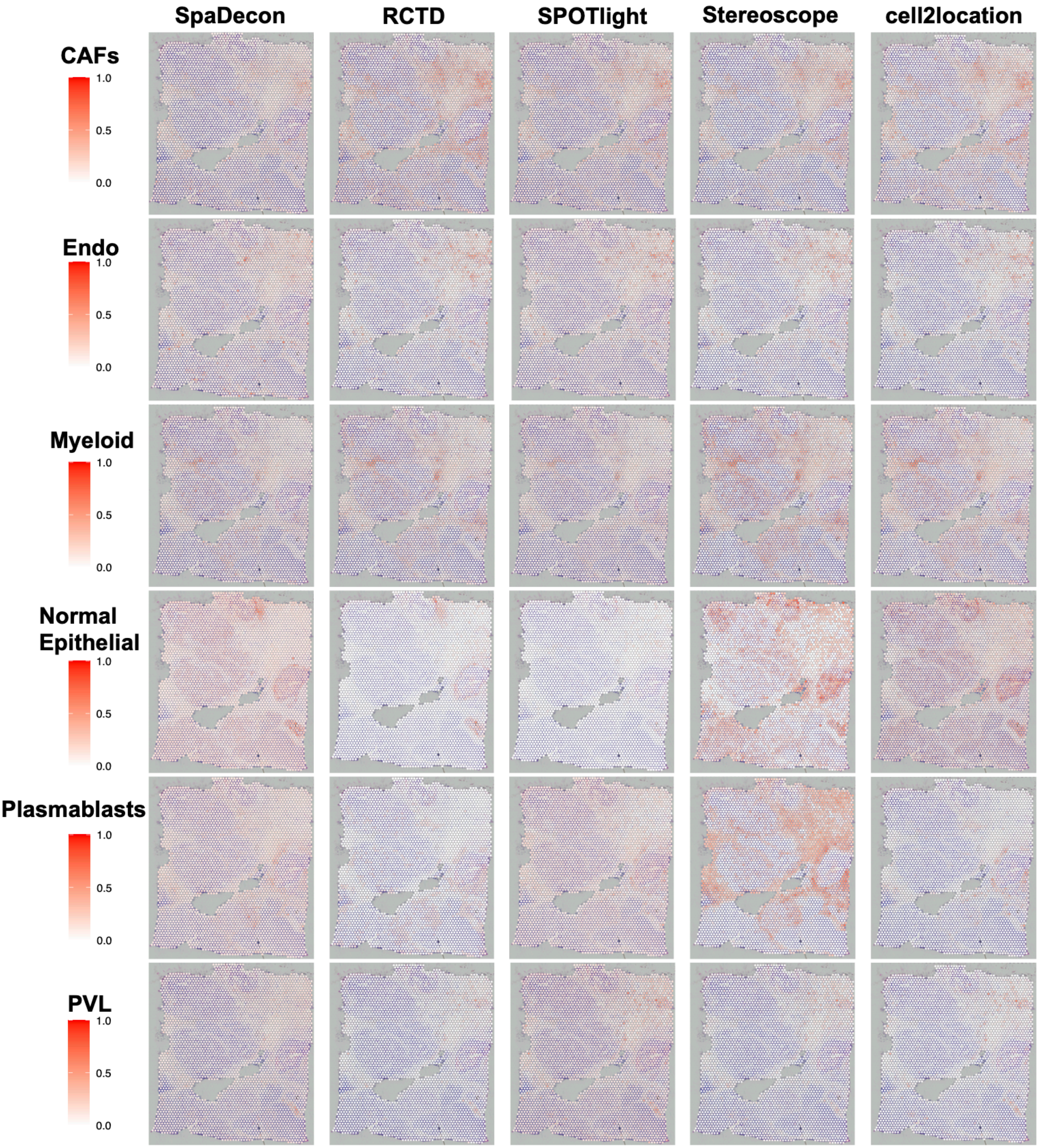

**Supplementary Figure 5.** For each method, heatmaps displaying the Jensen-Shannon Divergence (JSD) between the proportions estimated using the first scRNA-seq subset as reference and those estimated using each following scRNA-seq subset as reference for cell-type deconvolution of the 10X Visium breast cancer dataset.

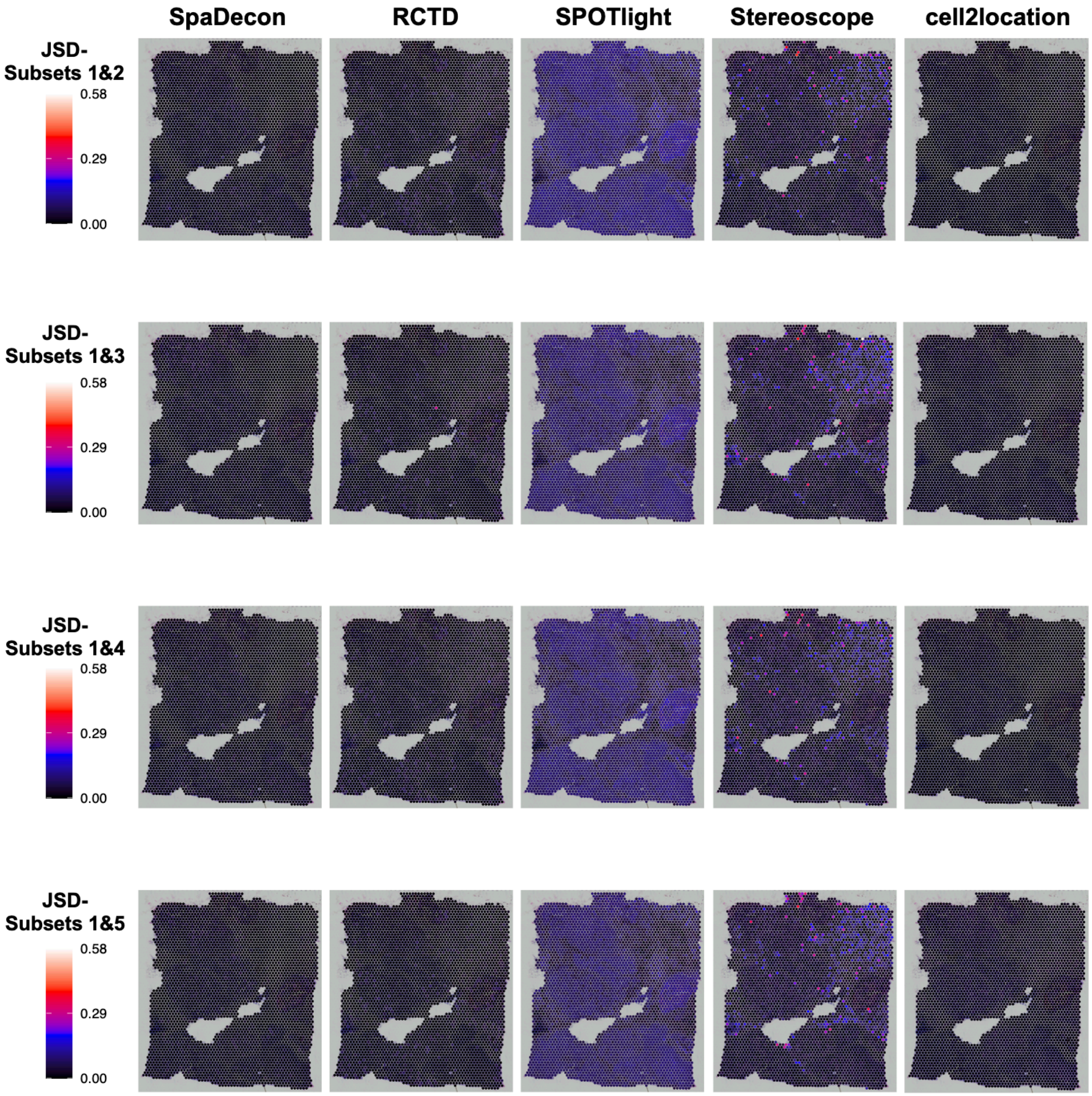

**Supplementary Figure 6. 10X Visium breast cancer benchmark evaluations. a,** Boxplot showing the mean squared error between the benchmark and estimated proportions across all cell types for each method. **b,** Boxplot showing the Jensen-Shannon divergence between the benchmark and estimated proportions across all spots for each method. **c,** Heatmaps showing the Jensen-Shannon divergence between the benchmark and estimated proportions at each spot in the pseudo-SRT dataset for each method.

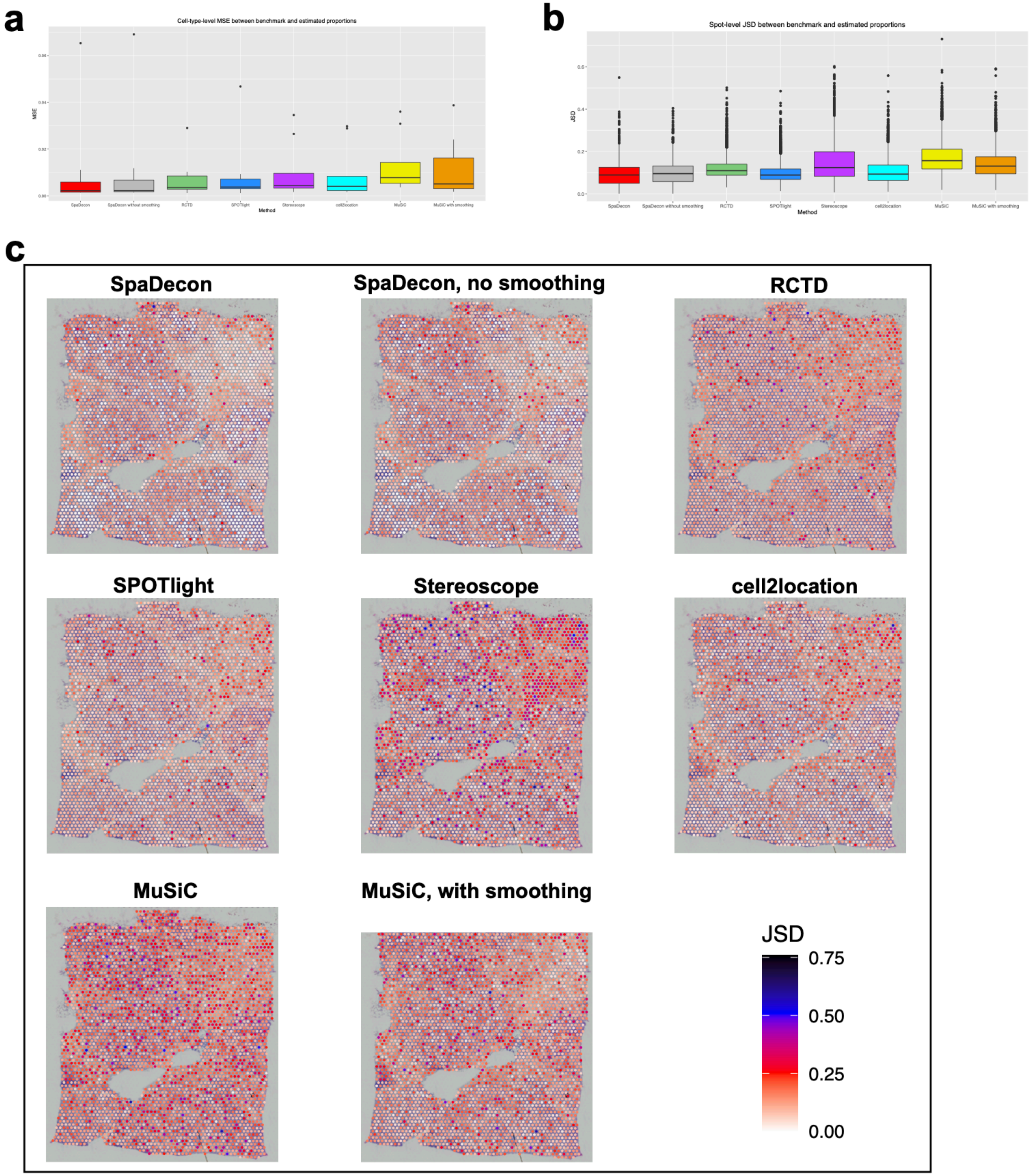

**Supplementary Figure 7.** For each method, heatmaps displaying the estimated distributions of cell types across the stage III cutaneous malignant melanoma ST dataset. Each spot is colored according to the proportion of a given cell type in that spot as estimated by a given method. See Supplementary Table 3 for full names of cell types.

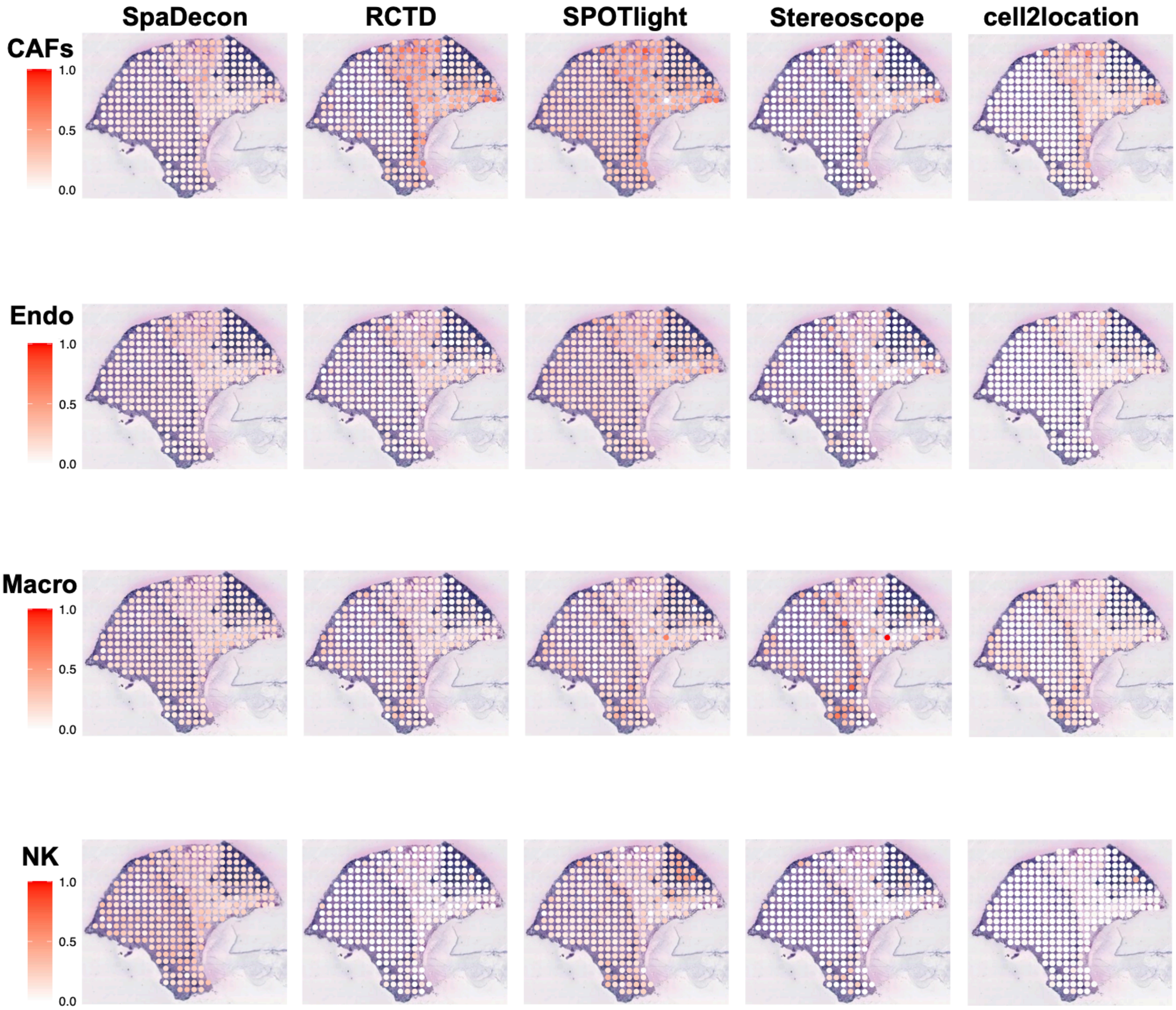

**Supplementary Figure 8. Stage III cutaneous malignant melanoma ST benchmark evaluations. a,** Boxplot showing the mean squared error between the benchmark and estimated proportions across all cell types for each method. **b,** Boxplot showing the Jensen-Shannon divergence between the benchmark and estimated proportions across all spots for each method. **c,** Heatmaps showing the Jensen-Shannon divergence between the benchmark and estimated proportions at each spot in the pseudo-SRT dataset for each method.

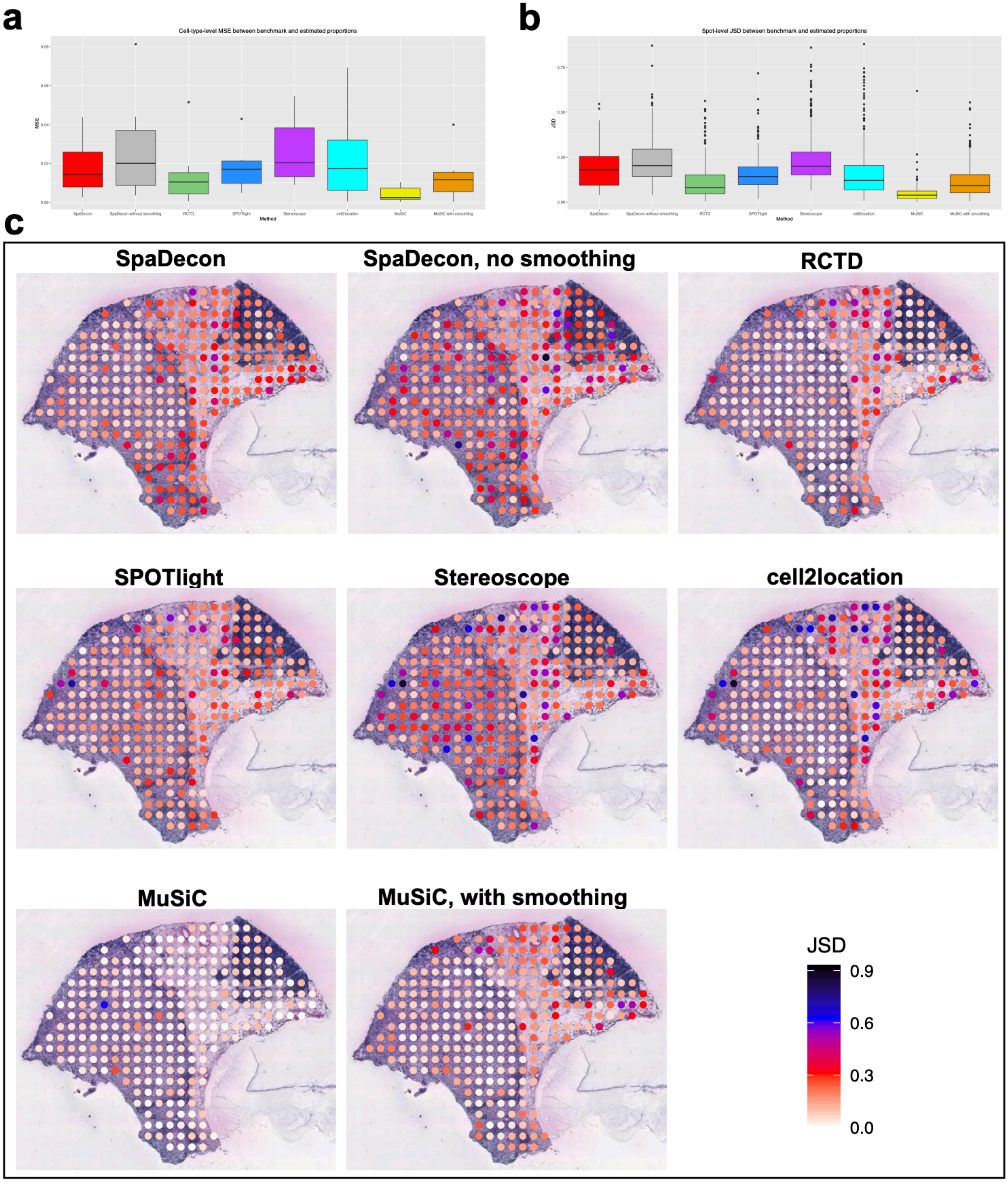

**Supplementary Figure 9.** For each method, heatmaps displaying the estimated distributions of cell types across the pancreatic ductal adenocarcinoma ST dataset. Each spot is colored according to the proportion of a given cell type in that spot as estimated by a given method. See Supplementary Table 3 for full names of cell types.

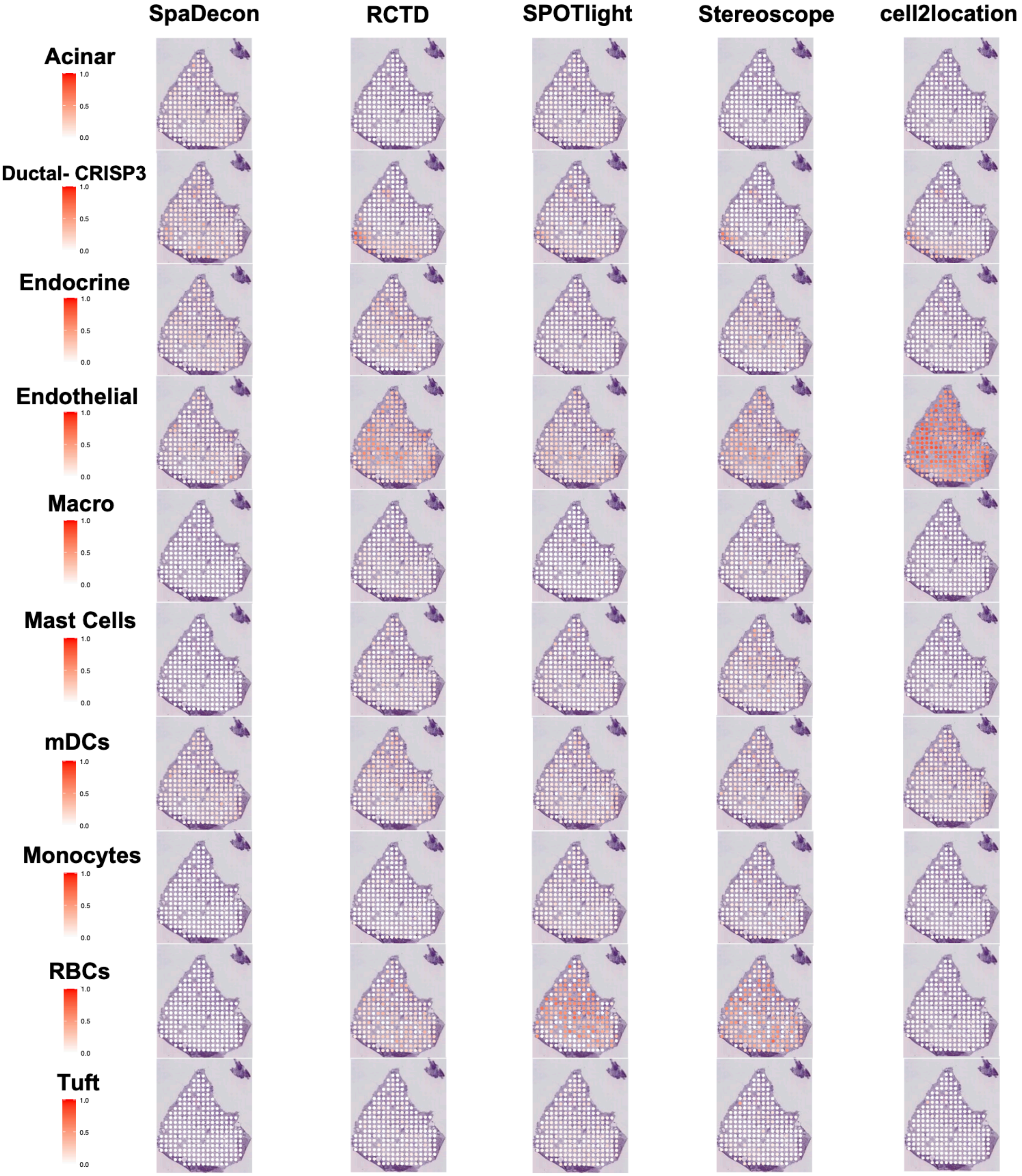

**Supplementary Figure 10.** Plot detailing the memory usage of SpaDecon when analyzing the 10X Visium breast cancer dataset. Memory was measured every 0.1 seconds and the maximum memory usage was approximately 5.06 gigabytes.

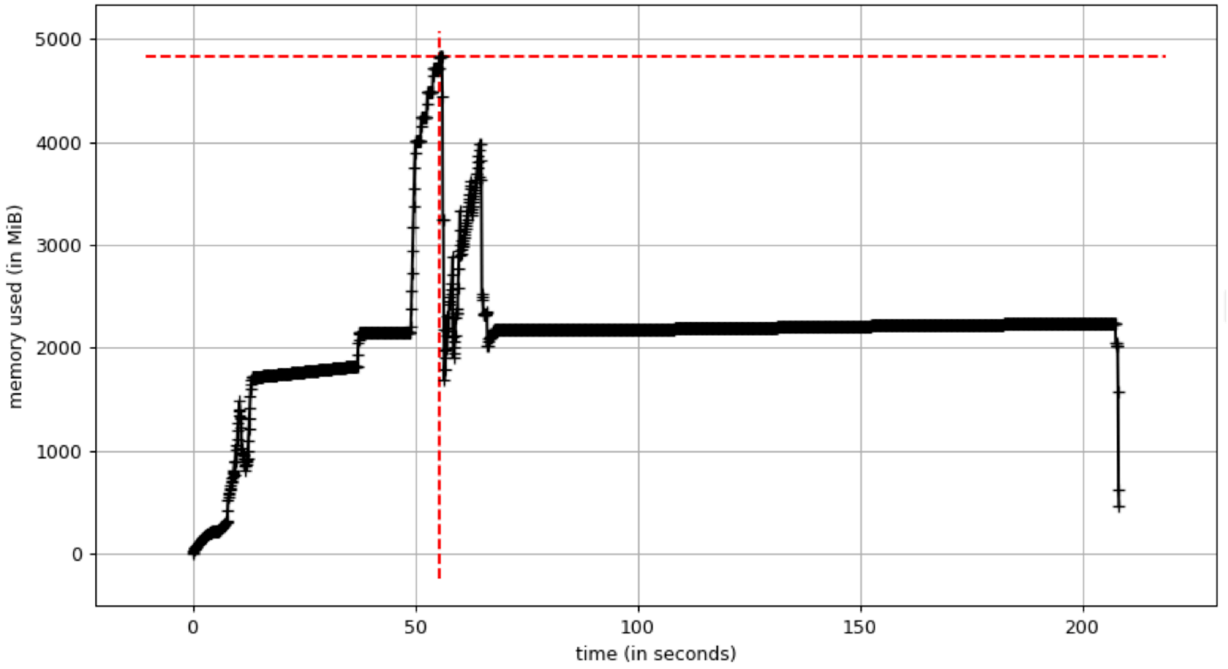
